# Copy number variation shapes structural genomic diversity associated with ecological adaptation in the wild tomato *Solanum chilense*

**DOI:** 10.1101/2023.07.21.549819

**Authors:** Kai Wei, Remco Stam, Aurélien Tellier, Gustavo A Silva-Arias

**Author notes:** Corresponding authors: Kai Wei, Aurélien Tellier Gustavo A. Silva-Arias.

## Abstract

Copy Number Variation (CNV) is a prevalent type of variation affecting large genomic regions which contributes to both genetic diversity and ecological adaptation in plants. The target genes involved in adaptation through CNV in tomato and its wild relatives remain unexplored at the population level. Therefore, we characterized the CNV landscape of *Solanum chilense*, a wild tomato species adapted to dry habitats, using whole-genome short-read data of 35 individuals from seven populations. We identified 212,207 CNVs, including 160,926 deletions and 51,281 duplications. We found a higher number of CNVs in diverging populations occupying stressful habitats. CNVs and single nucleotide polymorphisms analyses concordantly revealed the known species’ population structure, underscoring the impact of historical demographic and recent colonization events shaping genome-wide CNVs. Furthermore, we identified 3,539 candidate genes with highly divergent CNV profiles across populations. Interestingly, these genes are functionally associated with response to abiotic stress and linked to multiple pathways of flowering time regulation. Gene CNVs in *S. chilense* exhibit two evolutionary trends: gene loss in ancestral lineages distributed in central and southern coast populations and gene gain in the most recent diverged lineage from the southern highland region. Environmental association of the CNVs ultimately linked the dynamics of gene copy number to six climatic variables. It suggests that natural selection has likely shaped CNV patterns in stress-response genes promoting the colonization of contrasting habitats. Our findings provide insights into the role of CNV underlying adaptation during recent range expansion.

## Introduction

Copy number variation (CNV) is the primary type of structural variation (SV) caused by genomic rearrangements, which mainly includes deletion (DEL) and duplication (DUP) events resulting from the loss and gain of DNA segments (Feuk, et al. 2006; Żmieńko, et al. 2014). It is expected that CNV has a more significant impact on gene function than single-nucleotide polymorphisms (SNPs) because it covers more base-pairs (Shaikh, et al. 2009; Hämälä, et al. 2021) and has a higher per-locus mutation rate than SNPs (Lupski 2007). CNV is recognized as an essential driver of genomic divergence and local adaptation (Rinker, et al. 2019; Hämälä, et al. 2021; Marszalek-Zenczak, et al. 2023). Genome-wide studies confirm the importance of CNV in stress response and yield improvement in multiple plants, such as maize (Springer, et al. 2009), rice (Fuentes, et al. 2019; Qin, et al. 2021), and *Arabidopsis thaliana* (Zmienko, et al. 2020; Marszalek-Zenczak, et al. 2023). However, such studies have been conducted, so far, in selfing species and/or crops characterized by small effective population size (*N*_e_) and domestication bottlenecks (Alonso-Blanco, et al. 2016; Beissinger, et al. 2016; Brumlop, et al. 2019). Therefore, it is difficult in such species to disentangle the genome-wide effect of random evolutionary processes (genetic drift, chromosomal rearrangements, and demographic history) generating fast and extensive CNVs between populations from that of adaptive processes at specific loci (here positive selection underpinning environmental adaptation (Johri, et al. 2022). The dynamics of gene copy number indeed results from the population history and multiple events, including selection, migration and recombination (Sudmant, et al. 2015; Zhou, et al. 2019; Otto, et al. 2022; Antinucci, et al. 2023; Otto and Wiehe 2023). Indeed, the *N*_e_ of populations determines the amount of genetic diversity (SNPs or CNVs) available, the efficiency of positive and negative selection against genetic drift, and the effect of linked selection around sites under selection, thus being a major determinant of the genome architecture (Lynch and Walsh 2007). It is therefore more difficult to disentangle the effect of neutral processes from that of selection in small populations and/or populations with strong past demographic changes, for example following range expansions with strong bottlenecks (Johri, et al. 2022).

The tomato wild relative species *Solanum chilense* is an excellent model species to study the genetic basis of adaptive evolution when colonizing novel habitats (Böndel, et al. 2015; Stam, et al. 2019b; Wei, et al. 2023). Features such as outcrossing, gene flow, seed banks, and relatively mild bottlenecks during the colonization of new habitats result in high *N*_e_, as reflected by high nucleotide diversity and high recombination rates, meaning that this species has a high adaptive potential (Arunyawat, et al. 2007; Stam, et al. 2019b; Wei, et al. 2023). *S. chilense* occurs in southern Peru and northern Chile, from mesic to very arid habitats around the Atacama Desert, and is the southernmost distributed species in the tomato clade (Nakazato, et al. 2010). Moreover, within *S. chilense*, two lineages expanded southward during two independent colonization events (Böndel, et al. 2015; Stam, et al. 2019b; Raduski and Igić 2021; Wei, et al. 2023): one, early divergent towards the coastal part of northern Chile (hereafter the southern coast group, SC), and the other with a recent post-glacial divergence towards the high altitudes of the Chilean Andes (hereafter the southern highland group, SH) (Fig. 1A). The populations currently occurring in the southern coast and southern highland habitats have been shown to exhibit signatures of past positive selection for adaptation to cold, drought, light (photoperiod), heat and biotic stress (Xia, et al. 2010; Fischer, et al. 2011; Nosenko, et al. 2016; Böndel, et al. 2018; Stam, et al. 2019b; Wei, et al. 2023). These signatures of past adaptive selection suggest a genetic basis for the adaptation to novel habitats during the southward expansion of *S. chilense* populations towards arid areas around the Atacama desert (Wei, et al. 2024). Furthermore, these studies show that it is possible, to some extent, to disentangle in *S. chilense* the local footprints of strong positive selection (due to local adaptation) from the noise and variation in genome-wide polymorphism patterns due to neutral past demographic events. As advocated in (Johri, et al. 2022), we rely on providing orthogonal evidence from demographic inference and simulations guiding selective sweeps scans and correlation with climatic data. However, these studies revealed adaptive signatures based on scans for positive selection using solely SNP data: whether CNV can also contribute to adaptation to novel habitats in *S. chilense* is still unknown.

**Fig. 1.**
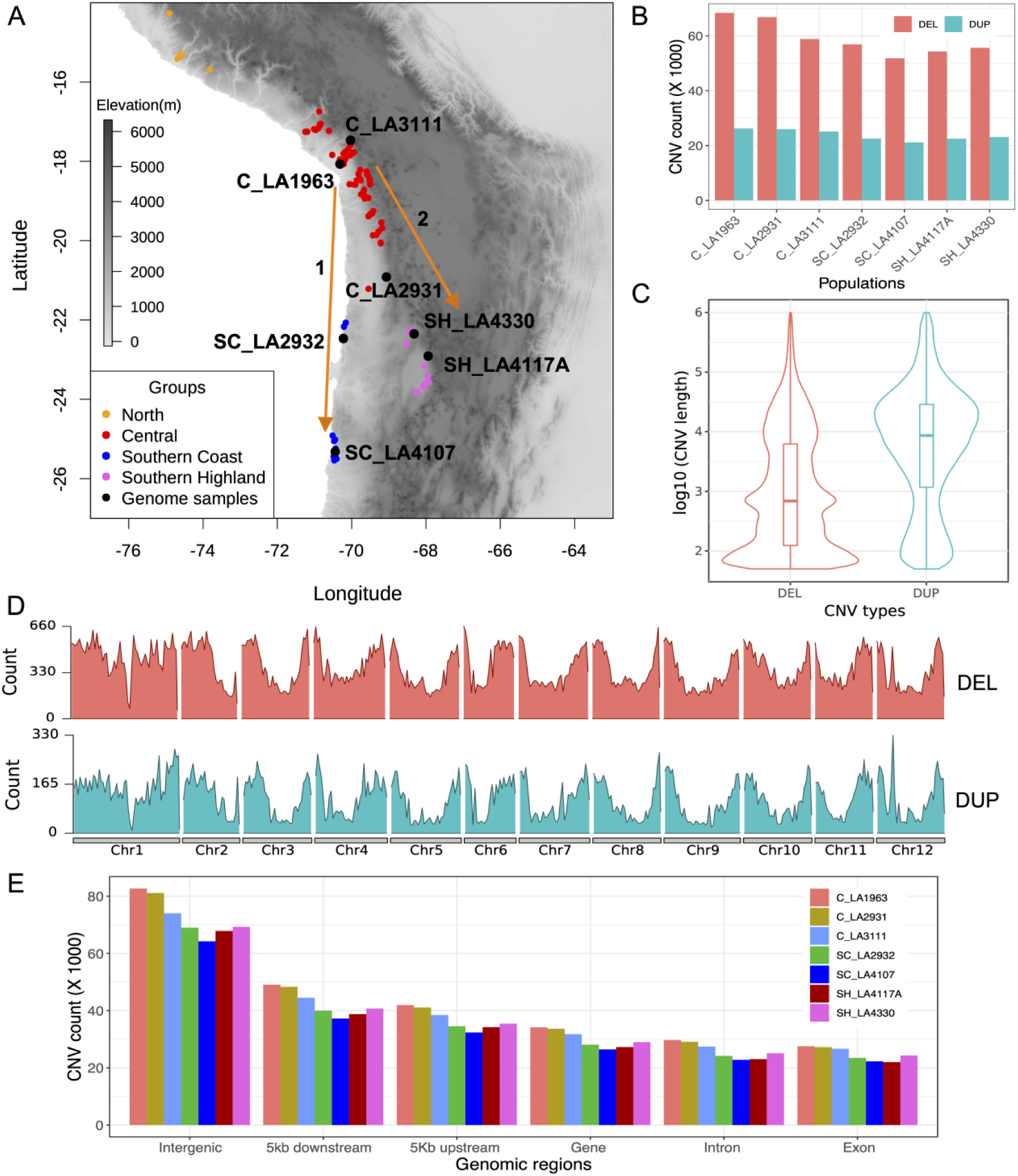
Overview of copy number variation detected in 35 *S. chilense* individuals. (A) The map with the distribution of all *S. chilense* populations at the Tomato Genetics Resource Center (TGRC), the seven *S. chilense* populations in this study (black circles), and the four population groups (circles with other colours). The two reconstructed southward colonization events, first to the southern coast and second to the southern highland (orange arrows). C: central; SH: southern highland; SC: southern coast. (B) The number of CNVs pooled across five individuals within each population. DEL: deletion; DUP: duplication. (C) The distribution of CNV size. (D) The CNV density along the genome is expressed as a count per 1Mb window. (E) The number of CNVs overlapping various genomic features for each population.

Reference genomes of several species of the tomato clade, including numerous cultivated tomato varieties, have been sequenced and assembled (Ranjan, et al. 2012; Sato, et al. 2012; Bolger, et al. 2014; Stam, et al. 2019a). Three tomato SV sets have recently been constructed based on a tomato-clade pangenome analysis to investigate the impact of genome rearrangements on gene expression and genomic diversity and provide new genomic resources for the improvement of tomato (Alonge, et al. 2020; Zhou, et al. 2022; Li, et al. 2023). These three studies compared cultivated tomato genomes with that of several wild tomato species, including an individual of the *S. chilense* population LA1969 (belonging to the central group; Fig. 1A). Interestingly, these studies showed that *S. chilense* exhibited the highest number of SV among all wild and cultivated tomato species, while the closely related wild tomato species *S. peruvianum* and *S. corneliomulleri* show only up to half of the number of SVs found in *S. chilense* (Li, et al. 2023). All these three species exhibit a similar recent proliferation of transposable elements (Li, et al. 2023). As *S. chilense* occurs in a wide range of environments, this species is of key importance for understanding the role of CNV in speciation and intraspecific diversification processes in the tomato clade. However, the studies mentioned above focused on the pangenome level across species (wild and cultivated), and an understanding of the role of CNV in local (ecological) adaptation is still lacking, especially for the adaptation to new arid habitats in southern populations in *S. chilense*.

In this work, we identified genome-wild CNVs and generated copy number (CN) for each gene based on genome-wide short-read sequencing data for 35 *S. chilense* individuals from seven populations (five diploid individuals per population) representing three different geographic habitats: three central (C) populations, two southern highland (SH) populations and two southern coast (SC) populations (Fig. 1A; Dataset S1). Based on these data, we first identified “candidate genes with highly differentiated CN” (CN-differentiated genes) across populations that are likely candidates associated with the inter-population differentiation in *S. chilense*. We then measured the evolutionary trend of expansion and contraction of gene CN based on candidate genes for a specified phylogenetic tree. Finally, we associated the dynamics of gene CN with climatic variables to provide evidence for environmental stresses driving CNV dynamics across populations. Our results suggest that CNV contributes to population adaptation to novel habitats in an outcrossing species with a large *N*_e_ and genetic diversity. We shed light on the importance of including an analysis of CNVs to complement genomic scans of recent positive selection based on SNPs.

## Results

### Summary of CNVs in the genome of *S.chilense* and validation of the pipeline

We identified a total of 212,207 CNVs (160,926 deletions and 51,281 duplications) using the combination of four CNV callers and the alignment of each of the 35 whole-genome sequencing datasets (Dataset S1) to the chromosome-level *S. chilense* reference genome (Silva-Arias, et al. 2025) (Fig. S1; Dataset S2). We found 73,014 to 94,621 CNVs per population (Fig. 1B; Table S1) and 31,923 to 46,579 CNVs per individual (Table S2). Although the number of deletions in all individuals and populations is much larger than the number of duplications (Fig. 1B; Fig. S1), the mean size of duplications (39,140 bp +/-104,577) is larger than that of deletions (14,052 bp +/-59,930) and exhibits a skewed distribution (Fig. 1C; Kolmogorov-Smirnov test, *P*=2.2e-16). We found 37% to 43% of the CNVs to be private to one individual in the three central populations. In comparison, only 12% to 14% of all CNVs are fixed in each of the three central populations (Fig. S2), *i.e.,* CNVs were observed in all five individuals of a given population. Southern populations (southern coast and southern highland) exhibited more fixed CNVs than the central populations, especially the two southern coast populations (25% in SC_LA2932 and 31% in SC_LA4107; Fig. S2).

Deletions and duplications were enriched at both ends of the chromosomes (Fig. 1D), consistent with previous studies (Alonge, et al. 2020; Hämälä, et al. 2021; Li, et al. 2023). Although most CNVs (76% to 79% per population) cover intergenic regions (Fig. 1E), about 35% to 38% of CNVs impacted coding sequences annotated in the *S. chilense* reference (some large CNVs were counted repeatedly due to covering multiple genes and intergenic regions). In addition, 45% and 50% of CNVs across populations overlapped with putative regulatory elements 5 kb upstream and 5 kb downstream of genes, respectively. As expected, 68% of deletions and 82% of duplications matched at least one transposable element annotated in the *S. chilense* genome, supporting that CNVs are predominately shaped by transposable elements (Fuentes, et al. 2019; Alonge, et al. 2020).

To confirm the validity of our pipeline, which assembled CNV detection from four tools specialized for short-read datasets, we simulated 1,000 deletions and 1,000 duplications with lengths ranging from 50 bp to 1 Mb based on 150 bp short reads (see methods). Our pipeline successfully detected approximately 90% of the simulated CNVs, and the false-positive rate was much lower than based on a single caller (Table S3). Our results, as well as previous claims, indicated that combining multiple callers can effectively improve the detection of CNVs based on short-read data (Kosugi, et al. 2019; Mahmoud, et al. 2019; Coutelier, et al. 2022).

### CNVs effectively capture the known species population structure

We compared the results of population structure analyses based on genome-wide SNPs and CNVs. The principal component analysis (PCA) based on the genotyped CNV dataset agreed with the clustering patterns from the genome-wide SNP dataset (Fig. 2A; Fig. S3A). Both analyses suggested a division of our samples into four genetic clusters that aligned with the geographic structure of the populations (best K = 4). The first principal component (PC1) separated the southern coast populations from inland (central and southern highland) populations, PC2 separated the southern coast subgroup into two genetic clusters (SC_LA2932 and SC_LA4107), and PC3 separated the inland populations into central and southern highland clusters (Fig. 2A; Fig. S3A). The ADMIXTURE analysis confirmed this result (Fig. 2B; Fig. S3B, with K = 4 exhibiting the lowest cross-validation error) and was consistent with the results from the SNP dataset (Fig. S3C).

**Fig. 2.**
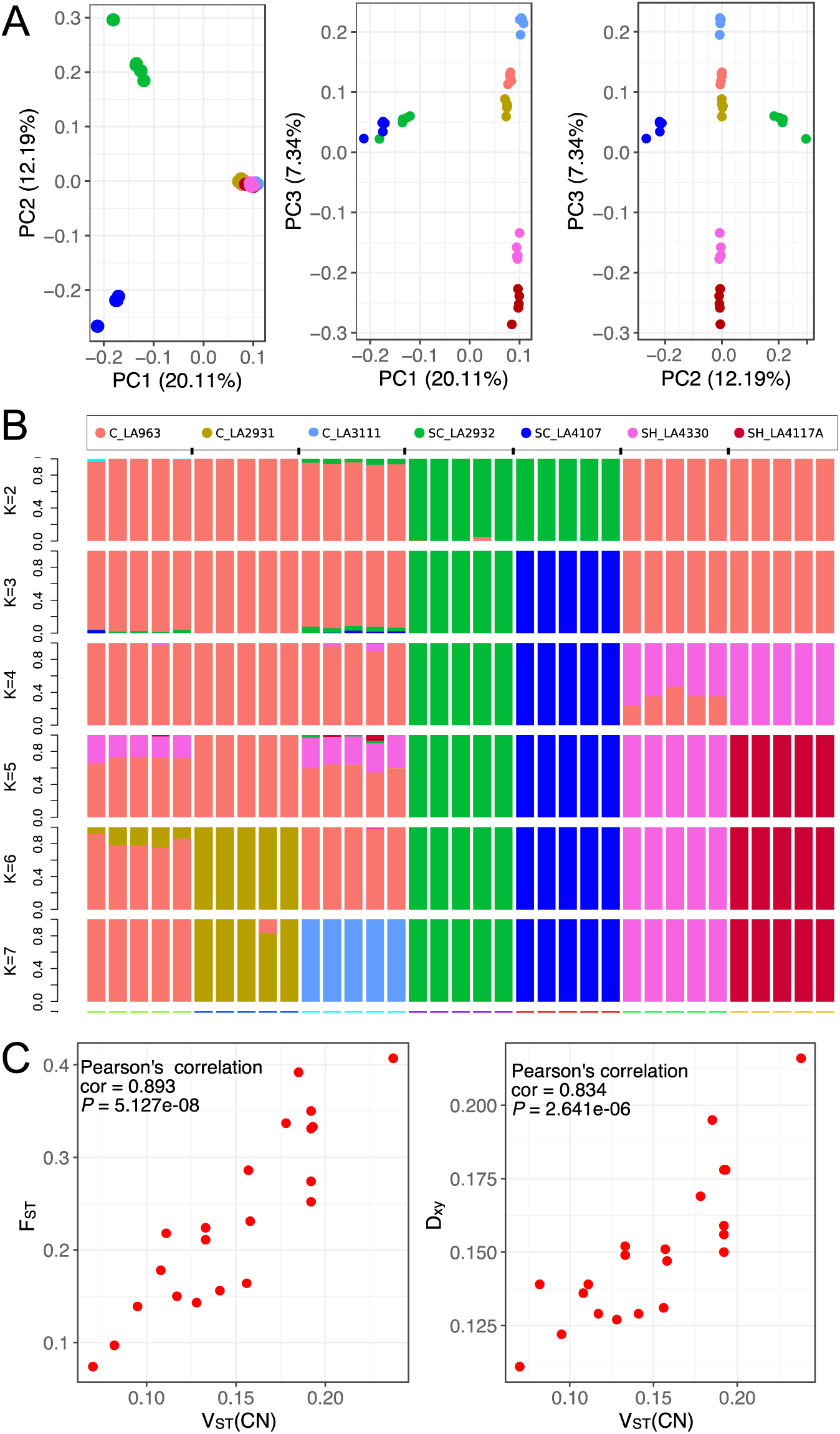
Population structure and differentiation analyses based on the genotyped CNVs. (A) Principal component analysis (PCA) based on the genotyped CNVs from 35 individuals from seven *S. chilense* populations. (B) Structure analysis based on genotyped CNVs and assuming between *K* = 2 and *K* = 7 subgroups (The best K value determined from the cross-validation error was 4; Fig. S3B). C: central; SH: southern highland; SC: southern coast. (C) The correlation between pairwise *F*_ST_/*D*_xy_ and pairwise *V*_ST_ indicates that CNVs support the known population differentiation.

We further explored the population differentiation using the *V*_ST_ statistics. This statistic is analogous to the classically used *F*_ST_ and *D*_xy_ statistics, but using CN values instead of allele frequencies (Redon, et al. 2006). The *V*_ST_ statistic ranges between 0 and 1, where 1 indicates that the populations are fully differentiated. We first computed the pairwise *V*_ST_ values along the whole genome in 1 kb windows using two CN quantitative measurements: Control-FREEC (*V*_ST_(CN)) and read depth (*V*_ST_(RD)) (Table S4). We found a highly significant positive correlation between these two estimators of the pairwise *V*_ST_ statistic (Pearson’s test, *P*=1.06e-07; Fig. S4A). In addition, all duplicated and lost fragments detected by Control-FREEC can be found in the CNV dataset obtained using the pipeline based on the four SV detection tools. Based on the pairwise *V*_ST_ statistics, we found similar structure patterns as in previous studies based on SNPs (Böndel, et al. 2015; Stam, et al. 2019b; Raduski and Igić 2021; Wei, et al. 2023), namely the high differentiation between southern coast and inland populations, especially between southern coast and southern highland populations (average *V*_ST_(RD)= 0.257 ± 0.039, average *V*_ST_(CN)= 0.198 ± 0.027; Table S4). As expected, both *V*_ST_ statistics (*V*_ST_(CN) and *V*_ST_(RD)) showed a highly significant positive correlation with *F*_ST_ and *D*_xy_ based on SNPs (Pearson’s test, *P* values see Fig. 2C; Fig. S4B to D).

### Differentiation of gene CN in different populations

To explore the role of natural selection in shaping CNV frequencies and distribution across populations, we also calculated global *V*_ST_ statistics (also with two methods, *V*_ST_(CN) and *V*_ST_(RD)) for each gene (39,245 genes in total). We aimed to capture candidate genes under divergent selective pressures by identifying genes with strong CN differentiation across all populations (Fig. S5). In total, we identified 3,539 candidate genes that present outlier CN differentiation across the seven populations (*i.e.,* genes with global *V*_ST_ greater than the top 95^th^ percentile of the 1,000 permuted *V*_ST_ values; Fig. S5; Table S5; Dataset S3) and 2,192 strongly CN-differentiated genes of these belong to the top 99^th^ percentile of the 1,000 permuted *V*_ST_ values (Fig. S5; Table S5). In Fig. S6, we show the distribution of deletions and duplications for these 3,539 candidate genes. Southern highland populations exhibited a comparatively large increase in gene gains (duplications) and a reduction in gene loss (deletions) relative to the other populations. In contrast, southern coast populations showed a comparatively high number of deletions relative to the high-altitude populations. In addition, southern highland and southern coast populations showed comparatively higher duplications than central populations (except C_LA3111). This may indicate that duplications play an important role during southward colonization.

We performed four PCA analyses based on the Control-FREEC–based CN values of 1) all annotated 23,911 genes with the mapped reads (Fig. S7A); 2) the 12,392 genes with *V*_ST_(CN)>0 (Fig. S7B); 3) the 3,539 differentiated gene set (observed *V*_ST_ values > 95% confidence interval cutoff in both gene CN estimate methods; Fig. 3A); and 4) the 2,192 strongly differentiated gene set (observed *V*_ST_ values > 99% confidence interval cutoff; Fig. S7C). In the PCA based on the 23,911 genes (Fig. S7A), all samples exhibited a cohesive grouping, except those from SC_LA4107. In the PCA based on the 12,392 genes with global *V*_ST_(CN) > 0 (Fig. S7B), two southern coast populations separated from the five inland populations (central and southern highland populations), suggesting a large difference in the CN range and composition between southern coast and inland populations. In the PCA based on the differentiated gene set (Fig. 3A; Fig. S7C), PC3 separated the southern highland populations from the central populations, consistent with the PCA based on the genotyped CNVs and SNPs (Fig. 2A; Fig. S2A). To rule out the effect of few outlier individuals on the PCA (Fig. S7A and B), we removed two outliers and found that the PCA results remain consistent (Fig. S8). Notably, however, that southern highland populations still showed ca. 20% of admixed ancestry coefficients with the central populations (Fig. 2B). These admixture signatures can reflect gene flow post-colonization of the southern habitats (between southern highland and central populations) or a very short divergence time. Consequently, similar polymorphisms in some parts of the genome were maintained between southern highland and central populations (Wei, et al. 2023). These results may indicate that the past demographic history of habitat colonization (and the resulting genetic drift) and gene flow are important evolutionary processes shaping both SNP and CNV frequencies within and between populations of *S. chilense*.

**Fig. 3.**
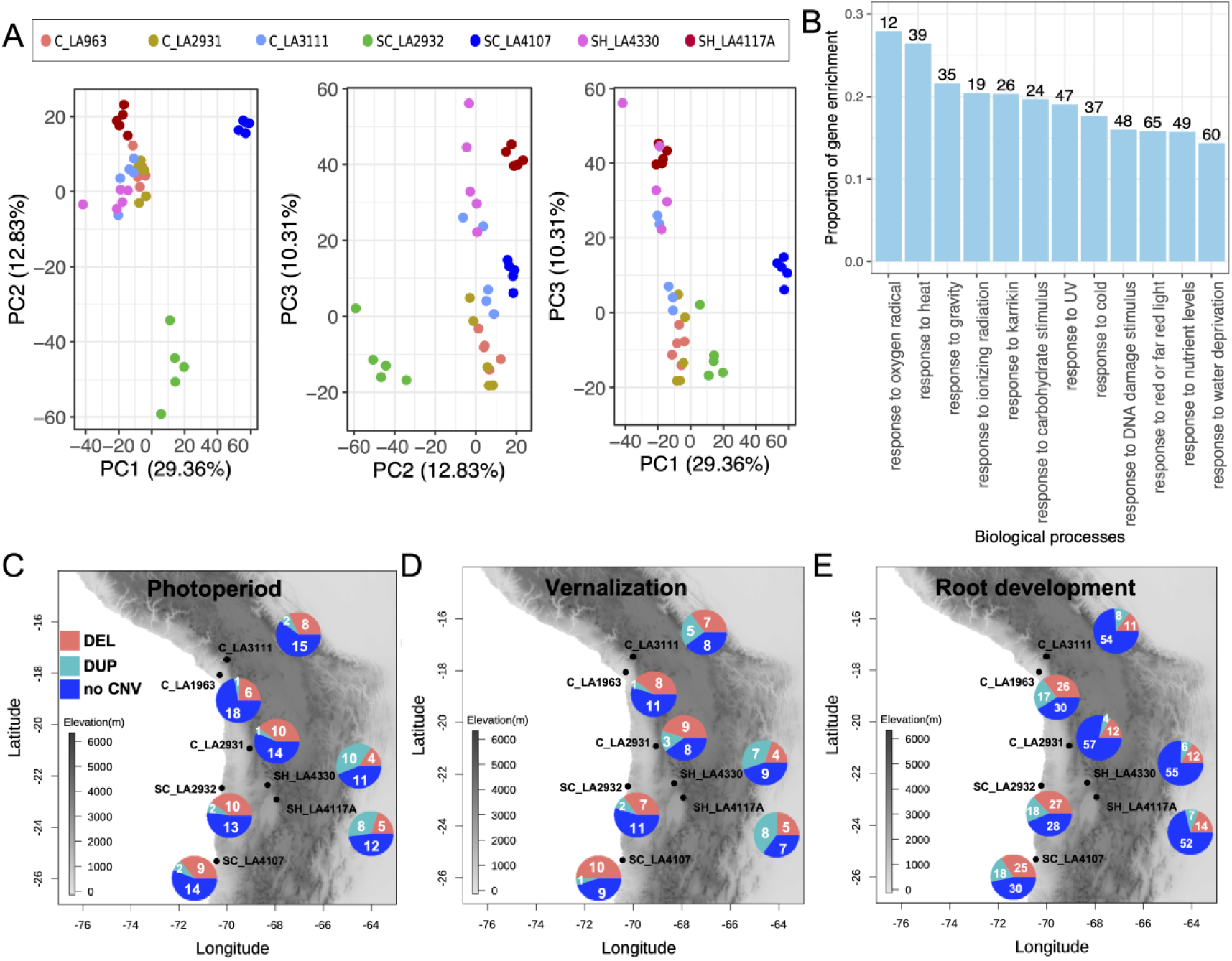
The CN-differentiated genes among seven populations are linked to response to multiple environmental stimuli. (A) A PCA based on the CN of 3,539 differentiated genes. C: central; SH: southern highland; SC: southern coast. (B) The proportions of CN-differentiated genes enriched in response to external stimuli/stresses (significantly enriched *P* < 0.05). The proportion of gene enrichment is defined as the number of genes enriched in one GO category divided by the number of background genes in this category. The number on each bar represents the number of genes enriched in that GO category. (C) The proportions of 25 CN-differentiated genes involved in the photoperiod pathway to regulate flowering time overlapping with deletion, duplication or absence of CNV in the seven populations. (D) The proportions of 20 CN-differentiated genes involved in the vernalization pathway to regulate flowering time overlapping with deletion (DEL), duplication (DUP) or absence of CNV (no CNV) in the seven populations. (E) The proportions of 73 CN-differentiated genes involved in the root developmental process overlap with deletion, duplication, or absence of CNV in the seven populations. The pie charts in C, D and E denote the proportions of CN-differentiated genes overlapping with deletion, duplication or absence of CNV (see also Table S6). The numbers in the pie chart indicate the number of genes overlapping with deletion, duplication or absence of CNV.

### Copy number variation illuminates enriched abiotic stress response pathways in *S. chilense*

We performed functional enrichment analysis on the 3,539 CN-differentiated genes according to GO biological process categories (Dataset S4). We classified the significantly enriched GO categories (*P* < 0.05) into nine groups (Fig. S9A) enriched for 82 genes (cell wall organization) up to 580 genes (cellular metabolic process). Interestingly, 400 (11.30%) CN-differentiated genes were enriched for a response to stimulus/stress that can be linked to multiple environmental factors (Fig. S9A), for example response to drought (water deprivation; 14.35% with 60 genes), cold (17.62% with 37 genes), heat (26.43% with 39 genes), red/far red light (15.82% with 65 genes), or ultraviolet light (UV; 19.03% with 47 genes) (Fig. 3B). The enrichment for these stress responses supported multiple sources of evidence for adaptation at genes associated with responses to arid conditions along a steep altitudinal gradient in *S. chilense* (Fischer, et al. 2011; Nosenko, et al. 2016; Böndel, et al. 2018; Blanchard-Gros, et al. 2021; Wei, et al. 2023). For instance, multiple drought-(*HSF* and *DREB3*), cold-(*FAD7*), and light/cold-responsive genes (*FT*, *GI*, and *FLD*) were found to be involved in flowering regulatory processes (Dataset S5). These findings are consistent with previous studies suggesting that selection pressures may occur at point mutations as well as at CNVs (Ofria, et al. 2003; Tan, et al. 2017; Lye and Purugganan 2019).

We found 227 CN-differentiated genes associated with flowering (Fig. S9A and B), an important fitness trait underlying local adaptation in plant species (Srikanth and Schmid 2011). As a critical part of the transition from vegetative to reproductive growth, flowering is influenced by multiple environmental conditions. Therefore, divergent flowering times related to local adaptation processes along the ecological gradient may be driven by CN-differentiated genes (Fig. S9C). We found 31 and 36 CN-differentiated genes linked to response to light and cold among the genes involved in flowering regulation (Fig. S9C), of which 25 and 20 genes were linked to photoperiod and vernalization pathways (Fig. S9B). The latter represent two regulatory flowering time pathways sensitive to the relative lengths of light-dark periods and low temperatures, respectively (Srikanth and Schmid 2011; Gaudinier and Blackman 2020). These genes showed a comparatively high overlap with duplications in southern highland populations (Fig. 3C and D; Fig. S10; Table S6). These genes included the potential homologs of floral integrator genes *FT* and *FD* (Liu, et al. 2008; Srikanth and Schmid 2011; Putterill and Varkonyi-Gasic 2016), putative homologs of *CRY2*, *GI*, and *ELF3* in the photoperiod pathway (Srikanth and Schmid 2011; Makita, et al. 2021), and a putative homolog of *AGL14* in the vernalization pathway (Hecht, et al. 2005; Pérez-Ruiz, et al. 2015). These candidate genes are well-known flowering time regulators in *A. thaliana* (Dataset S5). Note that these potential candidate genes related to flowering regulation were duplicated only in southern highland populations and exhibited either no CNVs or copy loss in central and southern coast populations (Fig. 3C and D; Table S6; t-test, *P* < 0.05). These findings indicate that genes with CN gains may promote colonization and adaptation in the southern highland habitats by regulating flowering time via the photoperiod and vernalization pathways (Wei, et al. 2023). Previous studies on several plant species have shown that a duplication of these positively regulated genes determining flowering time increases their expression level thereby promoting flowering (Blackman, et al. 2010; Díaz, et al. 2012; Panchy, et al. 2016). This genomic finding was consistent with the phenology observed in glasshouse conditions, in which southern highland individuals consistently flower 5-10 days earlier than those from central populations. In addition, other potential flowering regulatory genes in the differentiated gene set were likely involved in flowering regulation via different pathways (Dataset S5), namely the putative homologs of the genes *FY* and *FLD* (Srikanth and Schmid 2011; Cheng, et al. 2017; Bao, et al. 2020). The *FLD* gene showed an increased copy number in all populations (Dataset S5).

We identified 60 drought-responsive CN-differentiated genes associated with direct responses to water deprivation (Fig. 3B), encompassing duplicated homologs of *ABI4* and *AFP1* in the abscisic acid (ABA) pathway, along with a putative *WRKY33* transcription factor homolog with varying CN across populations (Dataset S5). These genes were validated as drought stress-responsive in *A. thaliana* and crops (Xiao, et al. 2021; Liu, et al. 2022; Luo, et al. 2022), including *WRKY33*, which is linked to temperature stress in tomato (Guo, et al. 2022). Furthermore, eleven CN-differentiated genes also belong to the drought-response metabolism co-expression network we previously found to be over-expressed under drought compared to well-watered conditions (Fig. S11; t-test, *P* = 2.68e-05) (Wei, et al. 2024), which corroborates their role in adaptive responses. Interestingly, we found similar numbers of deletion and duplication genes associated with water deprivation response across all populations (Fig. S9D; Table S6), suggesting a species-wide adaptation process in *S. chilense* through alterations in a metabolic gene network.

Our previous SNP study linked root development genes to putative local adaptation processes (primarily in response to extreme drought) in three low-altitude populations, including SC_LA2932, SC_LA4107 and C_LA1963 (Wei, et al. 2023). Accordingly, we also found 73 CN-differentiated genes involved in root development. These genes showed more CNVs in these three low-altitude populations (SC_LA2932, SC_LA4107, C_LA1963) than in high-altitude populations (C_LA2931, C_LA3111, SH_LA4117A, SH_LA4330) (Fig. 3E; Table S6; t-test, *P* < 0.05). This further indicated that root development may be an important strategy for adaptation to low-altitude environments.

### Gene expansion and contraction patterns show differences along altitudinal gradients

Our findings indicate that many CN-differentiated genes may be involved in adaptation to local habitats. To investigate the CN evolutionary trends of the 3,539 differentiated genes across populations, we performed an analysis of gene CN expansion (due to gene gain) and contraction (due to gene loss) across populations based on a phylogenetic tree derived from the inferred population genealogy (Fig. 4A). The CN of the differentiated genes was expanded (meaning there have been genes with CN gain) in the inland group with an expansion rate of 1.788 (Table 1). On the other hand, we found a gene reduction (meaning gene with CN loss) in the southern coast group with a contraction rate of –0.818. Within the inland group, the southern highland group exhibited CN gain (expansion rate of 0.416). In contrast, the central group showed CN losses (contraction rate of –0.767) three times higher than CN gains (Table 1). This likely indicates that gene CN of inland populations presents different evolutionary trends along the two evolutionary lineages. The two southern highland populations showed distinct CN expansion rates of 1.663 (SH_LA4117A) and 1.375 (SH_LA4330). In the central group, although the C_LA1963 and C_LA2931 displayed a trend of CN contraction, the C_LA3111 exhibited a similar rate of CN expansion (1.037) as the two southern highland populations (Table 1). The comparable CN expansion observed in the high-altitude populations (specifically, C_LA3111, SH_LA4330, and SH_LA4117A) may be attributed to three factors: the recent divergence of the southern highland group from the central group, the recent (re-)colonization of highland habitats following the glacial maximum (Wei, et al. 2023), and the ecological similarity of the habitats (Fig. 1A) which may also result in the duplication of a similar set of genes for C_LA3111 and the southern highland populations.

**Fig. 4.**
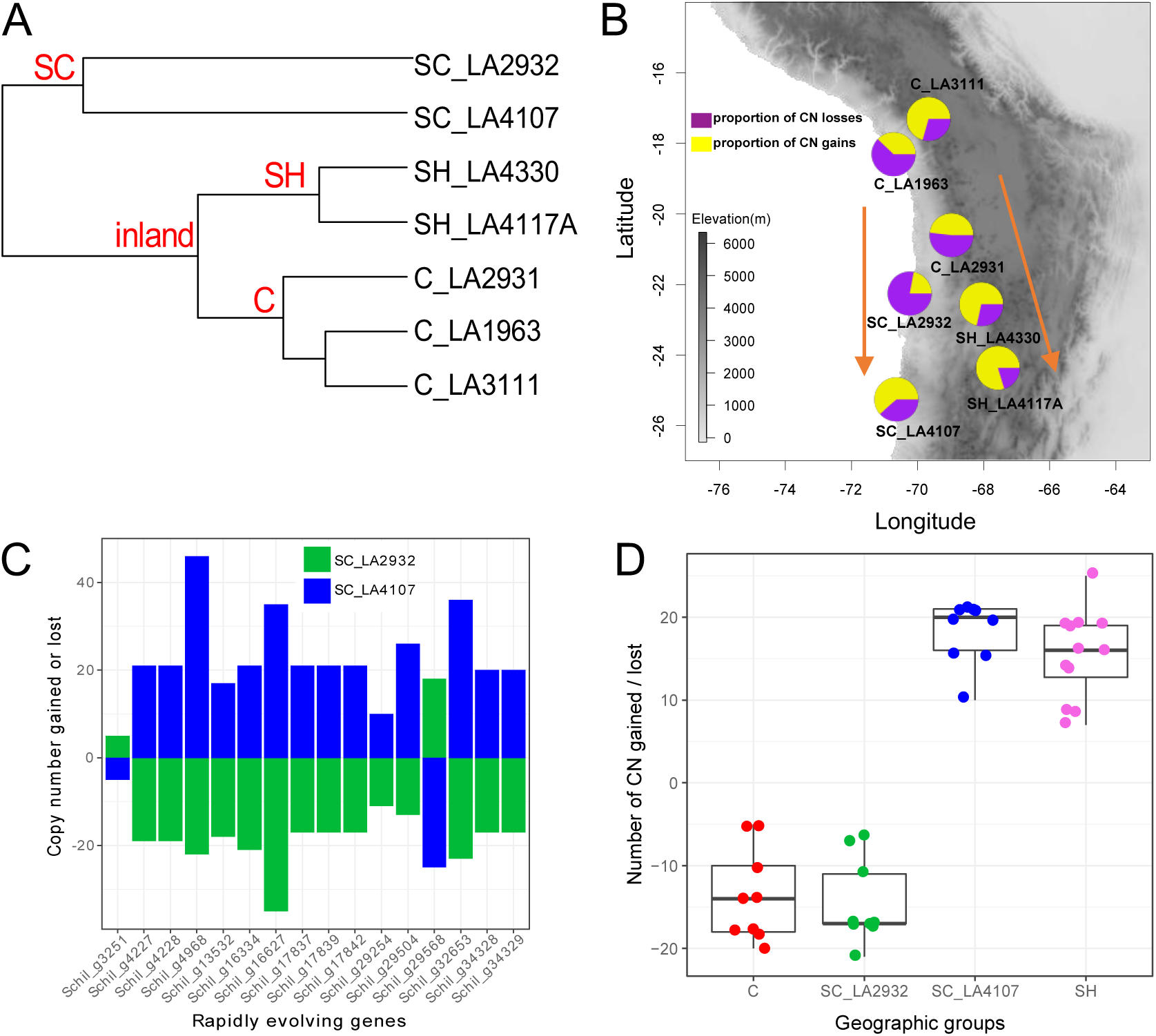
The expansion and contraction of CN-differentiated genes in different populations relative to the *S. chilense* reference genome. (A) The ultrametric phylogenetic tree used in gene expansion and contraction analysis (see Table 1). C: central; SH: southern highland; SC: southern coast. (B) The map and pie charts show the trends of gene CN loss and gain in the processes of two southward colonization events, first to the southern coast and second to the southern highland (orange arrows). The proportion of CN gains or losses for each population is defined as the number of CN gains or losses divided by the sum of the number of CN gains and losses. (C) The number of CN gains (positive values) or losses (negative values) for 16 rapidly evolving genes in two southern coast populations. (D) The number of CN gains and CN losses for rapidly evolving genes related to photosynthesis in four subgroups representing four different habitats (see also Table S8).

**Table 1.** The summary of gene expansion and contraction in different groups/populations based on an ultrametric tree.

| <sup>a</sup>Groups / Populations | Number of genes with CN expansion | Number of genes with CN contraction | Number of CN gained | Number of CN lost | <sup>b</sup>Rate of average expansion / contraction | <sup>c</sup>Number of rapidly evolving genes |
| --- | --- | --- | --- | --- | --- | --- |
| inland | 40 | 26 | 167 | 49 | 1.788 | 15 (+13/-2) |
| C | 163 | 695 | 355 | 1,013 | -0.767 | 20 (+5/-15) |
| SH | 527 | 525 | 1,143 | 705 | 0.416 | 37 (+32/-5) |
| SC | 48 | 359 | 106 | 439 | -0.818 | 9 (+2/-7) |
| C_LA1963 | 137 | 416 | 445 | 728 | -0.512 | 10 (+3/-7) |
| C_LA2931 | 212 | 458 | 815 | 878 | -0.094 | 15 (+3/-12) |
| C_LA3111 | 364 | 266 | 1,068 | 444 | 1.037 | 23 (+6/-15) |
| SH_LA4117A | 813 | 342 | 2,574 | 653 | 1.663 | 52 (+38/-14) |
| SH_LA4330 | 446 | 328 | 1,766 | 702 | 1.375 | 31 (+22/-9) |
| SC_LA2932 | 268 | 846 | 427 | 1,514 | -0.935 | 29 (+7/-22) |
| SC_LA4107 | 595 | 640 | 1,758 | 1,098 | 0.534 | 35 (+25/-10) |

<sup>a</sup>Groups and populations denote the branches in the ultrametric tree (Fig. 4A).
<sup>b</sup>Rate of average expansion / contraction = (Number of CN gained - Number of CN lost) / (Number of CN expanded genes + Number of CN contracted genes). Positive values indicate CN expansion and negative values indicate CN contraction.
<sup>c</sup>The rapidly evolving genes indicate significantly higher CN expansion or contraction ( $P < 0.05$ ) across the different groups/populations. Values outside parentheses represent the total number of the rapidly evolving genes. Positive values in parentheses denote the number of significantly expanded genes and negative values denote the number of significantly contracted genes (see also Dataset S6).

Interestingly, opposite results were observed between the two southern coast populations. Gene CN appeared to have contracted in SC_LA2932 (contraction rate of –0.935), while expansion occurred in SC_LA4107 (expansion rate of 0.534; Table 1) for the 3,539 differentiated genes. This follows our previous observation that the two southern coast populations showed a high degree of differentiation, possibly resulting from a long time of evolution in isolation and environmental differentiation. These results are also consistent with the population structure (Fig. 2) and may reflect the old southernmost colonization of the coastal habitats and the recent colonization of the highlands (Stam, et al. 2019b; Wei, et al. 2023).

Overall, the copy numbers of these potentially adaptively differentiated genes show an expansion (CN gain) in the two previously elucidated southward colonization events (Fig. 4B). Considering that the reference genome was assembled from population C_LA3111, which probably does not represent the ancestral state of the species. We also performed the same CN-expansion and contraction analysis using gene CN data calculated from the reference genome of *S. pennellii* (Table S7), a drought-adapted wild tomato species. We found consistent results, except for a slight decrease in the proportion of CN gains using the reference genome of *S. pennellii* in C_LA3111 (Fig. S12) compared to using the *S. chilense* reference (Fig. 4B).

We identified 155 “rapidly evolving genes” that exhibited higher CN expansion or contraction (see Methods) across the different groups/populations from 3,539 differentiated genes based on the reference genome of *S. chilense* (Table 1; Dataset S6). The 155 rapidly evolving genes also supported the population clusters in the PCA (Fig. S13), but C_LA3111 appeared closer to the southern highland populations than the other central populations. The highest number of such rapidly evolving genes were found in the southern highland populations (91 genes), including 71 significant CN expanded genes with GO enriched for photosynthesis (light reaction), long-day photoperiodism (flowering), and response to UV light and cold. We also observed 20 rapidly evolving genes primarily associated with developmental and metabolic processes. We also found 56 genes with rapidly evolving CN in the central populations (Table 1; Dataset S6), 75% of which exhibited a significant trend of CN contraction.

Among the 51 rapidly evolving genes in the two southern coast populations, 16 genes showed opposite CN trends in the phylogeny: a significant contraction in SC_LA2932 versus an expansion in SC_LA4107 (Fig. 4C). These genes included few homologs of photosystem subunits (i.e., *psb*B and *pet*D) mainly involved in photosynthesis (Dataset S5) and may underpin the high genetic differentiation at the CNV level between the two southern coast populations. In addition, the same CN rapidly evolving genes enriched for photosynthesis (light reaction) GO categories were also found in central and southern highland groups (Fig. 4D; Table S8). These potentially photosynthetic gene families appeared to have been contracting (CN loss) in the central group and SC_LA2932 but expanding (CN gain) in the southern highland group and SC_LA4107 (Fig. 4D; Table S8), suggesting that changes in the photosynthetic pathway may also be an important adaptive strategy across the different habitats in *S. chilense*.

### CN-differentiated genes are associated with climatic variation along the altitudinal gradient

To further explore CNV as the potential genetic basis of an adaptive response to abiotic factors, we conducted two genome-environment associations (GEA) analyses between the gene CN and 37 climate variables (Dataset S7).

We first implemented a redundancy analysis (RDA) to identify climate variables significantly associated with CN-differentiated genes across the seven populations. Three climatic variables (Bio7, Bio8 and Bio19) were observed to correlate with CN changes in the RDA based on 12,391 genes with global V_ST_(CN) > 0 (Fig. 14A). The first three RDA axes retained only 22.62% of the putative adaptive gene CNV and only weakly distinguished between inland and southern coast populations (Permutation test, *P* < 0.001; Fig. S14A to C). In the RDA based on the 3,539 CN-differentiated genes, 52.11% of the variance in CN can be explained by six climate variables (explanatory variables; sum of proportions in Fig. 5C) from five significant RDA axes (Permutation test, *P* < 0.001; Fig. 5A and 5C; Fig. S14D). These climatic variables were significantly correlated with the different populations (Mantel test, *P* < 0.05; Fig. 5B). In concordance with the PCA (Fig. 2A), the two main ordination axes did cluster the seven populations into four groups corresponding to the main geographical habitats (central, southern highland and two southern coast habitats). RDA axis 1 (RDA1) was correlated with the annual temperature range (Bio7) and potential evapotranspiration during the driest period (PETDriestQuarter). This axis represented the differentiation between the southern coast and inland populations (Fig. 5A and B). RDA axis 2 (RDA2) reflected the differentiation between two southern coast populations by mean temperature of the wettest quarter (Bio8). RDA2 also summarized a climatic gradient differentiating the low altitude (C_LA1963) and highland populations, which was mainly driven by solar radiation (ann_Rmean) and potential evapotranspiration (annualPET and PETColdestQuarter) (Fig. 5A and B). These six climatic variables were primarily associated with the colonization of southern highland and southern coast populations (Fig. 5B). The proportions of gene CN differentiation explained by these six climatic variables ranged from 0.02 (annualPET) to 0.136 (PETColdestQuarter) (Fig. 5C), in which PETColdestQuarter and PETDriestQuarter (0.121) exhibited the highest importance and correlated with inland and southern coast populations, respectively (Fig. 5A to C). Moreover, temperature changes (Bio7 and Bio8) also explained about 20.8% of the gene CN differentiation (Fig. 5C). Solar radiation (ann_Rmean) was a specific variable correlated with high altitude populations and explained 3.6% of gene CN differentiation (Fig. 5A to C). A consistent RDA model was obtained using the 2,192 strongly CN-differentiated genes (Fig. S14E to G). Finally, no significantly associated climate variables and RDA axes (Permutation test, *P* < 0.001) were obtained in the RDA applied on the 20,372 non-CN differentiated genes (Fig. S14H). This may corroborate that the CN-differentiated genes respond to external environmental stimuli in *S. chilense*.

**Fig. 5.**
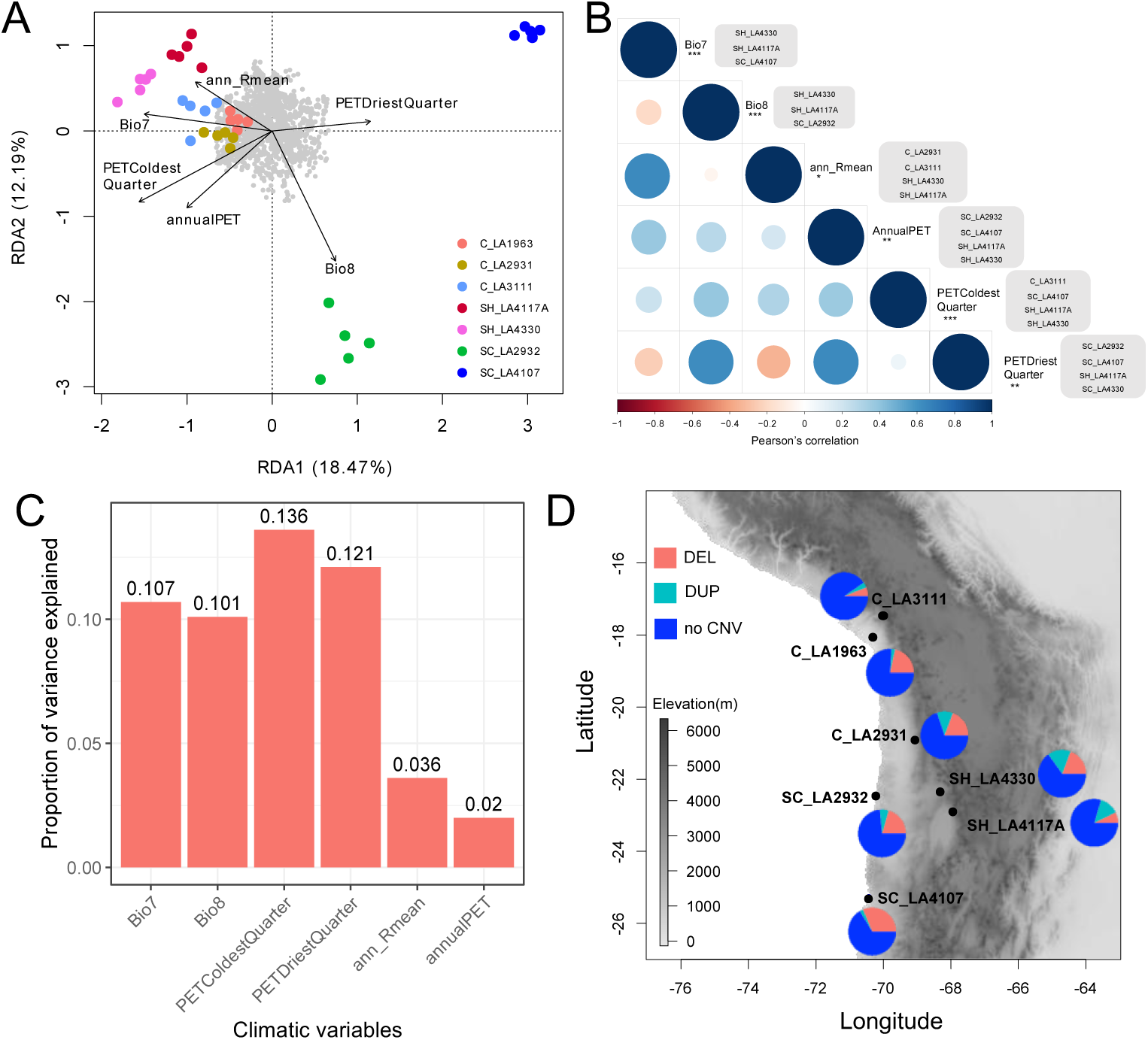
Genome-Environment Association (GEA) analysis between the gene CN and the climatic data of the different habitats. (A) Redundancy analysis (RDA) ordination biplot illustrating the association between the climatic variables (Dataset S7), individuals, and 3,539 differentiated gene CN. In the RDA, arrows indicate the direction of the climatic variables associated with the different populations, and the projection of arrows onto an ordination axis shows the correlation with that axis. The grey points denote the CN-differentiated genes. C: central; SH: southern highland; SC: southern coast. (B) The correlations between six overrepresented climate variables and populations, respectively. The bubble chart shows correlations between six climate variables. The asterisks (*) indicate the levels of significance of the climate variables for the RDA model (Permutation test; * *P* < 0.05, ** *P* < 0.01, *** *P* < 0.0001). The grey boxes to the right of the climatic variables show the populations significantly associated with that climatic variable (Mantel test, *P* < 0.05). (C) The proportion of variance explained by six overrepresented climate variables in the RDA model. (D) 34 CN-differentiated genes associated with both temperature annual range (Bio7) and solar radiation (ann_Rmean) in seven populations. The pie charts denote the proportions of CN-differentiated genes with deletion (DEL), duplication (DUP) or absence of CNV (see also Table S9).

We subsequently searched for candidate genes that may be associated with the six overrepresented climate variables using latent factor mixed models (LFMM) (Fig. S15A) (Frichot, et al. 2013; Caye, et al. 2019). Here, we performed an association analysis between the climatic variables and 3,539 highly CN-differentiated genes (not all genes). We identified 312 CN-differentiated genes significantly associated with the six climatic variables (z-test; calibrated *P* < 0.01; Fig. S15 B; Dataset S8). The PCA based on the CN of these 312 candidate genes displayed population clustering consistent with the one found in the RDA model (Fig. S16A; Fig. 5A), supporting that the six climate variables reflected gene CN changes across the species distribution. Among these 312 candidates, we found 217 genes to be significantly associated with three Potential Evapotranspiration (PET) climate variables (annualPET, PETDriestQuarter, and PETColdestQuarter), of which 98 genes were shared in at least two PET variables (Fig. S15B). Indeed, PET was the primary variable reflecting the drought status of the habitat. We noted that these PET-associated CN-differentiated genes were found across all populations (Fig. S16B and S16C), and were mainly GO-enriched in metabolic and root development processes. This is consistent with previous genomic and transcriptomic analyses showing that metabolic pathways and root development are important responses to drought stress (Wei, et al. 2023; Wei, et al. 2024). This result confirmed that drought tolerance is likely the main environmental pressure driving CN evolution across the population distribution of *S. chilense*. Furthermore, 69% (34 out of 49) of the genes associated with Bio7 were also observed to be correlated with ann_Rmean (Fig. S15B), which is likely a consequence of the correlation between Bio7 and ann_Rmean (Fig. 5B; Pearson’s correlation = 0.50). These genes were mainly duplicated in the southern highland populations and lost in the southern coast populations (Fig. 5D; Table S9). This result likely reflects that cold and high solar radiation are challenging conditions in southern highland populations (Dataset S7). Multiple duplicated genes associated with solar radiation (ann_Rmean) were enriched for a response to UV light in high-altitude populations, such as (likely) homologs of UV-B receptor *ARI12*, and DNA repair gene *REV1* (Dataset S5) (Tossi, et al. 2019; Thompson and Cortez 2020). In addition, we also found a few CN-differentiated genes, such as putative homologs of *CPD* (Dataset S5), which are related to pigment (anthocyanins) accumulation and were statistically associated with solar radiation variables.

We finally observed that the number of duplicated genes associated with the six climatic variables in the southern coast and especially southern highland populations was much higher than in the central populations (Fig. S16B). The analysis of GO enrichment above showed that these duplicated genes are involved in response to environments, including light, drought, cold, UV, and photosynthesis, such as the likely homologs of the genes *FT*, *FD*, and *ABI4* and genes involved in the formation of photosystem subunits (Dataset S5). The number of candidate genes found as deletions was highly consistent with the RDA results (Fig. S16C; Fig. 5A). For example, a large number of deletions in genes significantly associated with Bio8 occurred in SC_LA2932 (27 genes; Fig. S16C), far more than in other populations. Consistently the RDA results showed that CNVs in SC_LA2932 were also predominantly associated with Bio8 (Fig. 5A). The analysis of GO enrichment showed that most lost genes are related to plant growth and development. The GEA analyses confirmed the adaptive relevance of gene CN expansion and contraction: (i) the CN-differentiated genes in the central group appeared mainly as contraction genes (deletions) while these appeared as expansion genes (duplications) in the southern highland populations; (ii) the gene CN changes were linked to the climatic variables and associated with colonization of novel habitats at the southern edge of the species distribution; and (iii) the expansion/contraction of gene CN in different populations and RDA model also matched the population structure.

## Discussion

In this study, we explored the role of genomic CNV in the ecological adaptations of *S. chilense*. A set of key genomic CNVs in *S. chilense* populations were found to be highly correlated with the species colonization process and environmental variables and thus were likely implicated in the adaptive differentiation between populations, probably because of their major impact on gene expression (Fuentes, et al. 2019; Rinker, et al. 2019; Alonge, et al. 2020; Hämälä, et al. 2021; Li, et al. 2023). This confirms that CNV has ubiquitous roles in adaptive processes in ecology and evolution (Ż mieńko, et al. 2014; Castagnone-Sereno, et al. 2019; Lauer and Gresham 2019; Mérot, et al. 2020). To better understand the genetic basis behind the fitness effect of CNV in natural populations, we analyzed whole-genome (short read) data for 35 *S. chilense* individuals from seven populations, which allowed us to identify genome-wide CNVs. Our CNV calling pipeline resolved hundreds of thousands of CNVs in *S. chilense*. The number of CNV for each population of *S. chilense* was similar to numbers found in the previous tomato clade CNV based on a pan-genome study that included a single sample of *S. chilense* (Li, et al. 2023). CNVs were abundant across all chromosomes and frequently resided within, or in close proximity to, genes in the *S. chilense* genome (Fig. 1). Widespread CNVs in *S. chilense* genome exhibited similar performance as SNPs for the inference of population structure and differentiation between populations (Fig. 2; Fig. S3). Based on the demographic model we developed previously (Wei, et al. 2023) as a neutral null model and the dynamic changes of gene CN in two southward colonization events, our results supported that most CNV is likely shaped by neutral processes (Silva-Arias, et al. 2025). However, a genome-wide perspective allowed us to identify CNV likely related to the adaptive divergence in recently colonized regions in response to abiotic stress.

We conservatively identified patterns of gene CN differentiation that are likely to represent footprints of adaptive divergence. CN differences of these genes across different populations reflected the neutral and divergent selection process between populations, demonstrating that CNV must be considered to fully understand how selection shapes genomic structural diversity and local adaptation. Overall, the evolutionary processes generating CNV diversity and divergence were dominated by the demographic history of *S. chilense*, namely two southward independent colonization events. Gene CN appears expanded in the southernmost SC_LA4107 and southern highland populations, which underwent recent colonization events and exhibited lower population sizes (Stam, et al. 2019b; Wei, et al. 2023), while gene CN revealed a trend of contraction in the central and SC_LA2932 populations (close to the species’ center of origin). Therefore, we estimated that CN expansion and contraction likely reflect and underpin selective events during the two southward colonization events. Conversely, some plant species exhibit adaptive evolution by gene loss, for example, adaptive gene loss has been associated with changes in pollinators in *Petunia axillaris* (Hoballah, et al. 2007), *Ipomoea quamoclit* (Zufall and Rausher 2004) and *A. thaliana* (Shimizu, et al. 2008). Our study suggests that adaptive gene loss may also occur in genes involved in plant growth and development in central populations, and genes involved in photosynthesis in central and SC_LA2932 populations (Fig. 4D). These findings confirm the critical role of gene loss in adaptive evolution. Changes in CN at photosynthetic genes underpin population differentiation between SC_LA2932 (gene loss) and SC_LA4107 (gene gain), two populations in two different habitats on the southern coast. CN differentiated genes were also enriched in response to multiple abiotic stresses, such as red/far red light, cold, UV, or drought. These response processes can directly affect plant reproduction and growth and regulate flowering regulatory processes (Fig. S9). These findings agree with our results based on SNPs showing that the reproductive cycle, namely the regulation of flowering time, may play a key role in adaptation to abiotic stress in *S. chilense* (Wei, et al. 2023).

The regulation of flowering time involved in response to light (photoperiod) and cold (vernalization) appear as key adaptive pathways for *S. chilense* populations to colonize southern habitats as suggested by the analysis of genome-wide SNPs (Wei, et al. 2023). Here, we obtained further candidate genes based on CNVs enriched for flowering regulatory pathways and response to changes in photoperiod and cold. These genes (putative *FT*, *FD*, *FLD* homologs) are duplicated in the southern highland populations (Fig. S10). Solar radiation is also a challenging condition for plants at high altitudes. Many CN-differentiated genes were enriched for a function in response to UV light (Fig. 3B; Dataset S4), including homologs of genes involved in the anthocyanin accumulation in response to UV light. In plants, anthocyanin accumulation can improve the tolerance for drought, cold, salt and biotic stresses (Kaur, et al. 2023), especially anthocyanins act as potent antioxidants which help in eliminating Reactive Oxygen Species (ROS) molecules and protect the DNA damage under UV radiations (Catola, et al. 2017; Fang, et al. 2019). This may indicate that the gene CNVs in anthocyanin accumulation pathway are important for adaptation in high altitude populations of *S. chilense*. This follows a previous ecological niche study which suggested that *S. chilense* populations are expanding to the habitats of high altitude (Wei, et al. 2023). More generally, the large number of gene losses in response to environmental stresses may indicate that the reduction of the genome size is a powerful evolutionary driver of adaptation (Albalat and Cañestro 2016; Helsen, et al. 2020; Monroe, et al. 2021). Further functional validation will help understand the molecular mechanisms through which copy number variation drives adaptive evolution in natural populations.

To provide further evidence for selection (versus the footprints of past demography), our RDA analysis ultimately linked the dynamics of gene CN across populations to six climatic variables (Fig. 5A and B), of which five climatic variables were consistent with previous RDA results based on SNPs (Wei, et al. 2023). Similar CNV-environmental interactions have been observed in *A. thaliana* (DeBolt 2010; Zmienko, et al. 2020), *S. lycopersicum* (Alonge, et al. 2020), *Theobroma cacao* (Hämälä, et al. 2021), and *Oryza sativa* (Fuentes, et al. 2019; Qin, et al. 2021). Our results also highlight that CNV likely plays an essential role in response to the environments and in the southward colonization in *S. chilense*. CNVs, especially duplications in southern highland populations exposed to typical high-altitude stresses, were enriched in genes with functions related to cold, change of photoperiod and solar radiation. The CN changes of differentiated genes in southern coast populations mainly correlated with drought stress, such as root development, cell homeostasis, or cell wall maintenance. Interestingly, gene CN differentiation related to photosynthesis provided evidence for the genetic underpinning of the adaptive differentiation between SC_LA2932 and SC_LA4107, representing two different coastal habitats (Fig. 1A and 4C). These differentiated genes revealed opposite CN evolutionary trends between the two southern coast populations. Indeed, we saw different habitats as SC_LA2932 grows in dry ravines (quebrada) in Lomas formations, whereas SC_LA4107 grows in extremely fine alluvial soil (with even some running water). Moreover, these chloroplast genes were detected in the nuclear genome, consistent with widespread events of organellar gene transfers to the nuclear genome in tomatoes (Pesaresi, et al. 2014; Lichtenstein, et al. 2016; Kim and Lee 2018). Since the chloroplast genome is much more conserved than the nuclear genome in plants, the transfer of chloroplast genes to the nuclear genome with CNVs likely facilitates the increase in genetic diversity at nuclear copies of chloroplast genes, influencing the ecological adaptability of *S. chilense* (Daniell, et al. 2016). These putative adaptive signatures related to photosynthesis were not found in previous studies based on genome scans of SNPs (Wei, et al. 2023). The three central populations showed mainly a trend towards gene loss and low correlation with climatic variables (Fig. 5A and B). This trend is consistent with the fact that GEA analyses based on current climatic data have limited statistical power to detect old adaptive selection signals, whether based on SNPs or CNVs, due to the occurrence of multiple historical confounding events such as genetic drift, migration, and recombination (De Mita, et al. 2013; Manel, et al. 2016). The two central populations (C_LA2931 and C_LA3111) found at high altitudes exhibit few adaptive duplication signatures, but some as possible responses to cold and solar radiation, similar to those observed for the southern highland populations (Stam, et al. 2019b; Wei, et al. 2023).

Finally, we would like to stress that our study likely underestimates the amount and importance of CNV in *S. chilense* as we do not possess long-read data for all populations and our measure of outlier CNVs using global *V*_ST_ are likely conservatives. First, the tests with simulations based on the short-read data showed that our pipeline based on four tools to recover CNVs was likely conservative, which means that we probably missed some CNVs. Second, there may be bias in finding footprints of selection when using seeds from accessions maintained and propagated at the Tomato Genetics Resource Center (TGRC; UC Davis, USA), as we discussed previously (Wei, et al. 2023). Third, we also point out that the detection of CN-differentiated genes by the global *V*_ST_ statistics might be inflated because it is hard to correct for multiple testing (especially without a neutral demographic model of CNV evolution). We refrain from using the pairwise *V*_ST_ values to search for CNVs under selection because the sample size per population remained rather low (five diploids), but with higher sample sizes, such comparison of global versus pairwise *V*_ST_ would pinpoint more precisely to the population in which CNVs may have been selected. The availability of a new reference genome (Silva-Arias, et al. 2025) and a small number of populations sequenced with long-read (Li, et al. 2023) do open the path to sequence wild populations with long-read sequencing and a complete assessment of the importance of CNV at abiotic stress genes in *S. chilense*. We highlight here that contrary to common practice in SNP analyses (Wei, et al. 2023 and recommendation in Johri, et al. 2022) there is no standard procedure for detecting CNVs under selection, and we used here a permutation method based on *V*_ST_ (see also Rinker, et al. (2019). Nonetheless, the *V*_ST_ measure, despite our randomization procedure, may be biased by low-frequency CNVs (as is known for *F*_ST_), and thus we used RDA to provide orthogonal evidence. Therefore, there is a need to develop new simulation and inference methods to study, infer and disentangle the neutral and selective processes driving gene duplication and deletion (Otto, et al. 2022; Otto and Wiehe 2023). These are much needed options to quantify/infer the neutral rates of gene duplication/deletion during the species southward expansion and local adaptation, and, thereby, develop robust statistical selection tests for CNVs. Fourth, instead of using the CN dataset of all genes to perform association analyses with climate variables, we used genes with high CN differentiation. The reason for this is that in the RDA analysis we did not obtain any associated climate variables when using CN dataset for all genes, indicating that the large number of genes with weak CN changes greatly reduced the resolution of the analysis (while low-frequency CNV are not picked up by the RDA analysis). This may confirm that the RDA complements our *V*_ST_ analysis and supports the footprints of selection (versus that of neutral processes) at our high CN-differentiated genes.

Despite being conservative regarding the importance of positive selection shaping the CNV diversity in *S. chilense*, our results reinforced the observation that CNV is an important contributor to adaptation across different ecological habitats (Żmieńko, et al. 2014; Rinker, et al. 2019; Hämälä, et al. 2021; Monroe, et al. 2021). The strong selective pressure imposed by the range expansion of *S. chilense* and the need to adapt to novel stressful habitats has shaped the genetic diversity at SNPs and CNVs. In agreement with previous studies, we suggest that natural selection acting on CNVs can reshape the genomic composition of populations and might form a basis for local adaptation (Iskow, et al. 2012; Żmieńko, et al. 2014; Rinker, et al. 2019; Hämälä, et al. 2021).

## Materials and Methods

### Sample collection and sequence read processing

The 35 *S. chilense* plants were grown in standard glasshouse conditions from seeds obtained from the Tomato Genetics Resource Center (TGRC, University of California, Davis, CA, USA). We sampled five diploid plants from accessions representing the three main geographic groups. We retrieved whole-genome short-read sequencing data from 35 specimens from seven populations of *S. chilense* (accession: C_LA1963, C_LA3111, C_LA2931, SH_LA4330, SH_LA4117A, SC_LA2932 and SC_LA4107; five diploid plants for each population) representing three main geographic groups and environments (Fig. 1A). The data are available on the European Nucleotide Achieve (ENA; BioProject accession no. PRJEB47577). We executed the same pipeline of read processing procedure as in our previous study (Wei, et al. 2023b), including quality trimming, mapping and SNP calling based on the reference genome of *S. chilense* (Silva-Arias, et al. 2025). The results of the sequencing and read mapping were documented in Dataset S1.

### Identification and genotyping of CNVs

To obtain high-confidence CNVs including deletions and duplications, we chose four software tools for SV detection based on an evaluation of SV detection tools by Kosugi et al. (2019). This study found that combining SV detection tools tends to give higher precision and that LUMPY (Layer, et al. 2014), Manta (Chen, et al. 2016), Wham (Kronenberg, et al. 2015) and DELLY (Rausch, et al. 2012) showed the best overall performance. These tools implement different calling algorithms that jointly draw information from patterns of read pairs, split reads, read depth, and de novo assembly.

For LUMPY v0.3.1, we first extracted the discordant paired-end reads with abnormal insertion size from mapped results using ‘*view*’ function of Samtools v1.7 (Wysoker, et al. 2009), and then we extracted the split-read alignments using the ‘extractSplitReads_BwaMem’ script. We used the ‘*sort*’ function of Samtools to sort the resulting BAM files. Next, we ran LUMPY using the mapped reads, discordant paired-end reads and split reads as inputs to detect CNVs. For CNV calling with DELLY v0.7.6, we chose the default parameters and converted the output file from bcf to vcf format using bcftools v1.9 (Danecek, et al. 2011; Danecek, et al. 2021). We also ran Manta v1.6 and Wham v1.8 using default parameters. For each individual, we merged the CNV sets obtained with these four tools using SURVIVOR v1.0.7 (Jeffares, et al. 2017). We set the minimum CNV length to 50 bp and the maximum CNV length to 1 Mb. Only CNVs of the same type (deletion or duplication) and same DNA strand (sense strand or antisense strand) detected by different tools were integrated. We retained CNVs that were called by at least two of the four tools.

We finally used the merged CNV set as input for SVTyper v0.7.0 to call breakpoint genotypes of the structural variants (Chiang, et al. 2015). The script we used for CNV calling, merging, and breakpoint estimation is available from our Gitlab repository: https://gitlab.lrz.de/population_genetics/s_chilense_cnv/-/blob/main/pipeline_of_CNV_calling_genotyping.

To assess the sensitivity and accuracy of our pipeline for CNV calling, we simulated short-read data with CNV using CNV-Sim v0.9.2 (https://github.com/NabaviLab/CNVSim). We simulated 1,000 duplication and 1,000 deletion regions ranging from 50 bp to 1 Mb based on 150 bp paired-reads. We then used our same pipeline to call CNVs based on this simulated short-read dataset (Table S5). The script implementing these CNV simulations is available from https://gitlab.lrz.de/population_genetics/s_chilense_cnv/-/blob/main/CNVs_simulation.

### Population structure analysis

We inferred the population structure using the whole-genome SNPs and genotyped CNVs. We performed the principal component analysis (PCA) to seek a summary of the clustering pattern among sampled genomes using GCTA v1.91.4 (Yang, et al. 2011). We first converted vcf format to plink format using VCFtools v1.17 (Danecek, et al. 2011), then converted plink format to a binary format using PLINK v1.9 (Purcell, et al. 2007) with parameters ‘--noweb –-make-bed’ to generate input of GCTA. We next performed the analysis of admixture using the program ADMIXTURE v1.3.0 (Alexander, et al. 2009). We assessed six scenarios (ranging from K = 2 to K = 7) for genetic clustering using the same input as the PCA analysis.

### Quantification of gene copy number

We employed two strategies to quantify gene copy number (CN). First, we used the read-depth based method implemented in Control-FREEC v11.6 to estimate the CN in 1 kb sliding windows across the entire genome (Boeva, et al. 2012). We used the following parameters in Control-FREEC: ploidy = 2, breakPointThreshold = 0.8, degree = 3, minExpectedGC = 0.3, maxExpectedGC = 0.55, and telocentromeric = 0. We then obtained gene CN from the Control-FREEC outputs and gene coordinates in the genome. However, some genes had more than one CN estimate. These events may be due to imperfect estimation of breakpoints using our window size. So, we calculated the average CN if one gene corresponded to multiple CN values.

We also employed an alternative strategy. We first extracted read depth using Mosdepth v0.3.2 (Pedersen and Quinlan 2018) in 1 kb sliding windows from BAM files, and then we calculated the read depth for each gene from gene coordinates. We used median read-depth values of all windows and genes as a normalizing factor to obtain the final window and gene CN estimate, respectively, and the formula reads as: CN = (read depth / median value) × 2. A factor of 2 stands for the species diploidy (Rinker, et al. 2019).

### Estimation of the population differentiation by CNVs

We calculated *V*_ST_ to estimate the population differentiation. The *V*_ST_ measurement, analogous to *F*_ST_, is applied to identify loci that differentiate by CN between populations (Redon, et al. 2006; Zhao and Gibbons 2018; Rinker, et al. 2019). *V*_ST_ is calculated by defining (*V*_T_ – *V*_S_)/*V*_T_, where *V*_T_ denotes the total variance and *V*_S_ denotes the average variance within each population, weighted by the sample size (five for all populations in this study). We first calculated, using a sliding window-based approach, pairwise *F*_ST_ and pairwise *V*_ST_ to compare the of measures of population differentiation by SNPs and CNVs. We calculated for each pair of populations the *F*_ST_ statistics using VCFtools over 1 kb sliding windows and *V*_ST_ based on CN of 1kb sliding window across the reference genome. Note that we calculated pairwise *V*_ST_ based on two different CN estimation strategies: using control-FREEC (*V*_ST_(CN)) and read depth (*V*_ST_(RD)).

### Identification of CNV candidate genes associated with the population differentiation

We identified candidate genes with significant CN differences between populations, the so-called CN-differentiated genes, using a global *V*_ST_ per each gene based on the gene CN (Zhao and Gibbons 2018; Rinker, et al. 2019). The per gene global *V*_ST_ calculation follows:

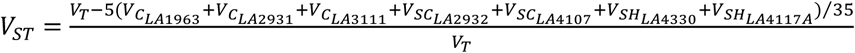

where *V*_T_ is the CN variance over all 35 individuals, *V*_pop_x_ is the CN variance for each respective population, five is the sample size for each respective (pop_x) population and 35 is the total sample size. An R script with the pipeline of gene *V*_ST_ calculations and identification of candidate genes is found on: https://gitlab.lrz.de/population_genetics/s_chilense_cnv/-/blob/main/VST.R. We performed permutation tests on the CN counts to identify which genes displayed the greatest degree of observed inter-population CN differentiation while controlling for sampling bias. Here, we randomly permuted gene CN of each gene for 35 individuals and calculated a new global *V*_ST_ for every permutation and every gene, respectively. We repeated 1,000 times the permutations to generate a random distribution of global *V*_ST_ values for each gene. We then selected candidate genes for which the observed global *V*_ST_ fell above the 95^th^ and 99^th^ percentile of the permuted global *V*_ST_ distribution. These candidate genes displayed strong intra-population CN homogeneity and high degrees of inter-population differentiation. Finally, genes were considered significant when observed *V*_ST_ values were above the maximum 95% (differentiated) or 99% (strongly differentiated) confidence interval cutoff in both gene CN estimation methods control-FREEC (*V*_ST_(CN)) and read depth (*V*_ST_(RD)) (the *V*_ST_ cutoff see Table S5).

### Gene ontology (GO) analysis

We first performed a BLAST (Camacho, et al. 2009) of our CN differentiated genes to the *A. thaliana* dataset TAIR10 (e-value cutoff was10^-6^). We selected the best matching entry (lowest e-value) as the target homologue for enrichment analysis. We performed a GO enrichment analysis using the *A. thaliana* annotation database as the background using the R package clusterProfiler (Yu, et al. 2012). When we determined the enriched GO terms, we used the Benjamini-Hochberg method (Benjamini and Hochberg 1995) to control the false discovery rate fixed at 0.05.

### Expansion and contraction of gene copy number

To gain insights into the changes of the gene CN size across populations in a way that accounts for phylogenetic history, we performed an analysis of gene CN expansion and contraction with the set of 3,359 differentiated genes using CAFE v4.2.1 (Han, et al. 2013). This program can estimate the evolution of the gene CN size based on a stochastic birth and death model. For a specified phylogenetic tree, and given the gene CN sizes in each individual, CAFE can calculate the global birth and death rate of gene CN. Then it infers the most likely gene CN sizes at all nodes in the tree and detects genes that have accelerated rates of CN gains and losses. It finally computes a *p-*value associated with each gene CN and identifies significant rapidly evolving genes with smallest *p*-values.

We first calculated the mean CN for 3,539 CN-differentiated genes for each population, respectively. We then constructed a population-based phylogenetic tree using SNPs by TreeMix v1.13 (Pickrell and Pritchard 2012), and then the ultrametric tree (Figure 4A) was generated based on ‘*force.ultrametric*’ function of phytools R package (Revell 2012). Finally, we analysed gene CN expansion and contraction in different groups. We first ran CAFE for genes with CN less than 100 to calculate an accurate *lambda* value (λ=0.00207 in this study), because genes with large CN can lead to non-informative parameter estimates. We then ran CAFE for genes with CN (gene copies) larger than 100 using the same *lambda* value calculated from genes with CN less than 100. We chose a significance threshold of 0.05 (*p*-value) when identifying rapidly evolving genes with an excess rate of evolution (expansion or contraction) in different groups/populations. The code we used to analyse CN expansion and contraction can be found on: https://gitlab.lrz.de/population_genetics/s_chilense_cnv/-/blob/main/run_cafe.sh.

### Association analysis between gene copy number and climatic variables

We obtained the environmental data, including 37 climatic variables, from two public databases, WorldClim2 (Fick and Hijmans 2017) and ENVIREM (Title and Bemmels 2018) (Dataset S7). To evaluate the relative contribution of the abiotic environment to explaining patterns of genetic variation, we first performed a redundancy analysis (RDA) to associate CN of 3,539 differentiated genes with climatic variables. We performed the RDA analysis using the *rda* function from the R package vegan (Forester, et al. 2018), modelling CN as a function of predictor variables and producing constrained axes and representative predictors (climatic variables). We assessed the multi-collinearity between representative predictors (climatic variables) using the variance inflation factor (VIF) and excluded all climatic variables with a VIF of 10 or above. We then calculated the significance of the RDA ordination axes using the *anova.cca* function (*P* < 0.001). The R script for the RDA analysis, including all steps and parameters, can be obtained at https://gitlab.lrz.de/population_genetics/s_chilense_cnv/-/blob/main/RDA.R.

We obtained six climatic variables significantly correlated with the changes of gene CN across populations from the RDA (Fig. 5A). To identify candidate genes associated with the climatic variables, we used LFMM2 (latent factor mixed models) to build a model between each gene and climatic variable based on the univariate test (Caye, et al. 2019). We used the *lfmm_ridge* function in the R package LFMM to obtain an object that contains the latent variable score matrix under the assumption of K = 4 latent factors (as evaluated from analysis of population structure) based on the CN of 3,539 differentiated genes and six representative climate variables (as obtained from RDA), respectively. Then, we performed association testing using the *lfmm_test* function. We finally used the method of Benjamini-Hochberg to calibrate *p*-values and set conservatively 0.01 as the significance threshold to obtain candidate genes associated with the climatic variables. The R script of LFMM we used is available on our Gitlab repository https://gitlab.lrz.de/population_genetics/s_chilense_cnv/-/blob/main/lfmm.R.

### Supplementary material

Supplementary data are available online at Molecular Biology and Evolution.

### Data Availability

Raw sequence data are available at the European Nucleotide Achieve (ENA) BioProject PRJEB47577. The resource of copy number variation identified in this study and custom scripts for conducting the analyses are available at our Gitlab at the following link: https://gitlab.lrz.de/population_genetics/s_chilense_cnv.

## Acknowledgements

KW was funded by the Chinese Scholarship Council. GAS-A was funded by the Technical University of Munich. KW acknowledges funding from Natural Science Foundation of Xinjiang Uygur Autonomous Region Grant Number: 2024D01C216 and “Tianchi Talents” introduction plan. AT acknowledges funding from DFG (Deutscheforschungsgemeinschaft) Grant Number: 317616126 (TE809/7-1). We thank the Tomato Genetics Resource Center (TGRC) of the University of California Davis for generously providing us with the seeds of the population included in this study, and the Greenhouses & Phytochambers Unit of the TUM Plant Technology Center in Dürnast for plant care.

## Competing interests

The authors have no conflicts of interest to declare.

## Author contributions

KW, GAS-A and AT planned and designed the study. RS and AT obtained the sequencing data. KW performed data analyses. KW wrote the first draft of the manuscript, and RS, GAS-A, and AT edited and improved the manuscript. All authors approved the final manuscript.

## Notes

### Competing Interest Statement

The authors have declared no competing interest.

### Summary of Updates

We have done some minor updates to clarify the statistical methods used to find CNV outliers, and improve at few places the writing.

