## Supplementary material for "Copy number variation shapes structural genomic diversity associated with ecological adaptation in the wild tomato *Solanum chilense*": SI information

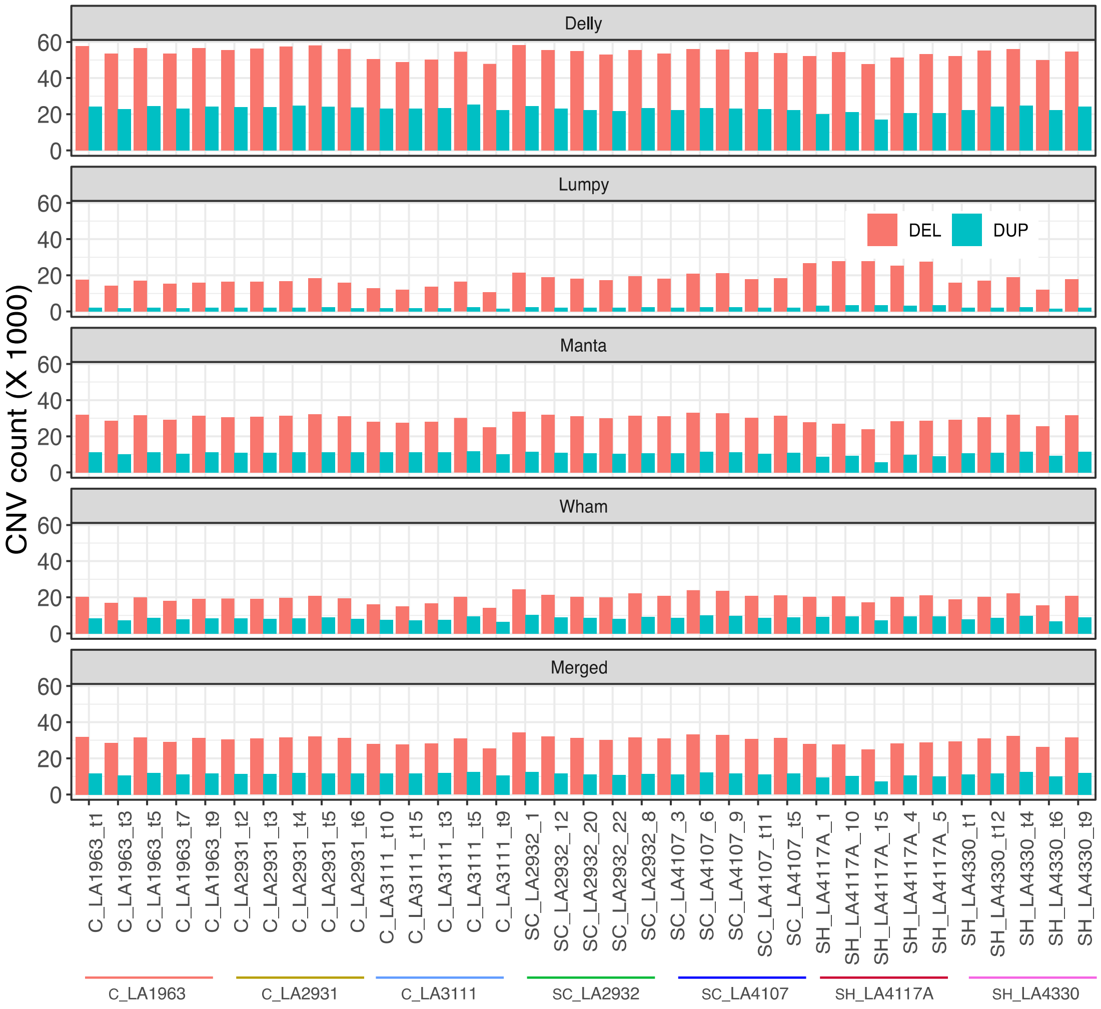


**Fig. S1.** The summary of deletion (DEL) and duplication (DUP) using four CNV callers in 35 individuals, and consensus result based on CNV calls supported by at least two callers. See also Dataset S2 and Table S2.

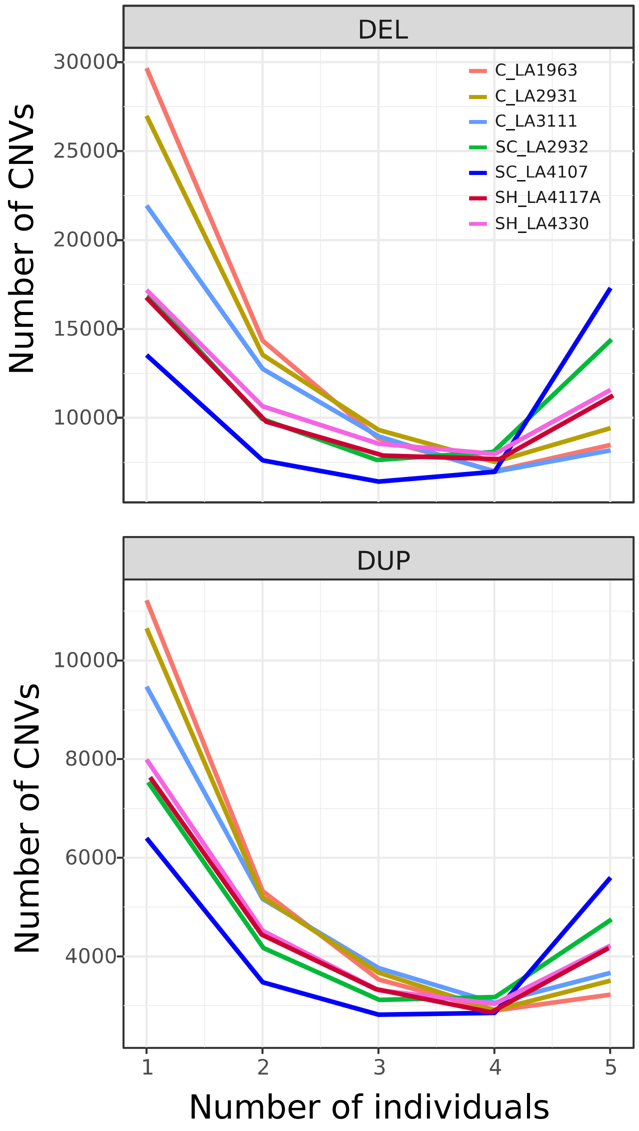


**Fig. S2.** The number of CNVs identified in 1, 2, 3, 4 or 5 individuals in each population, respectively. DEL: deletion; DUP: duplication.

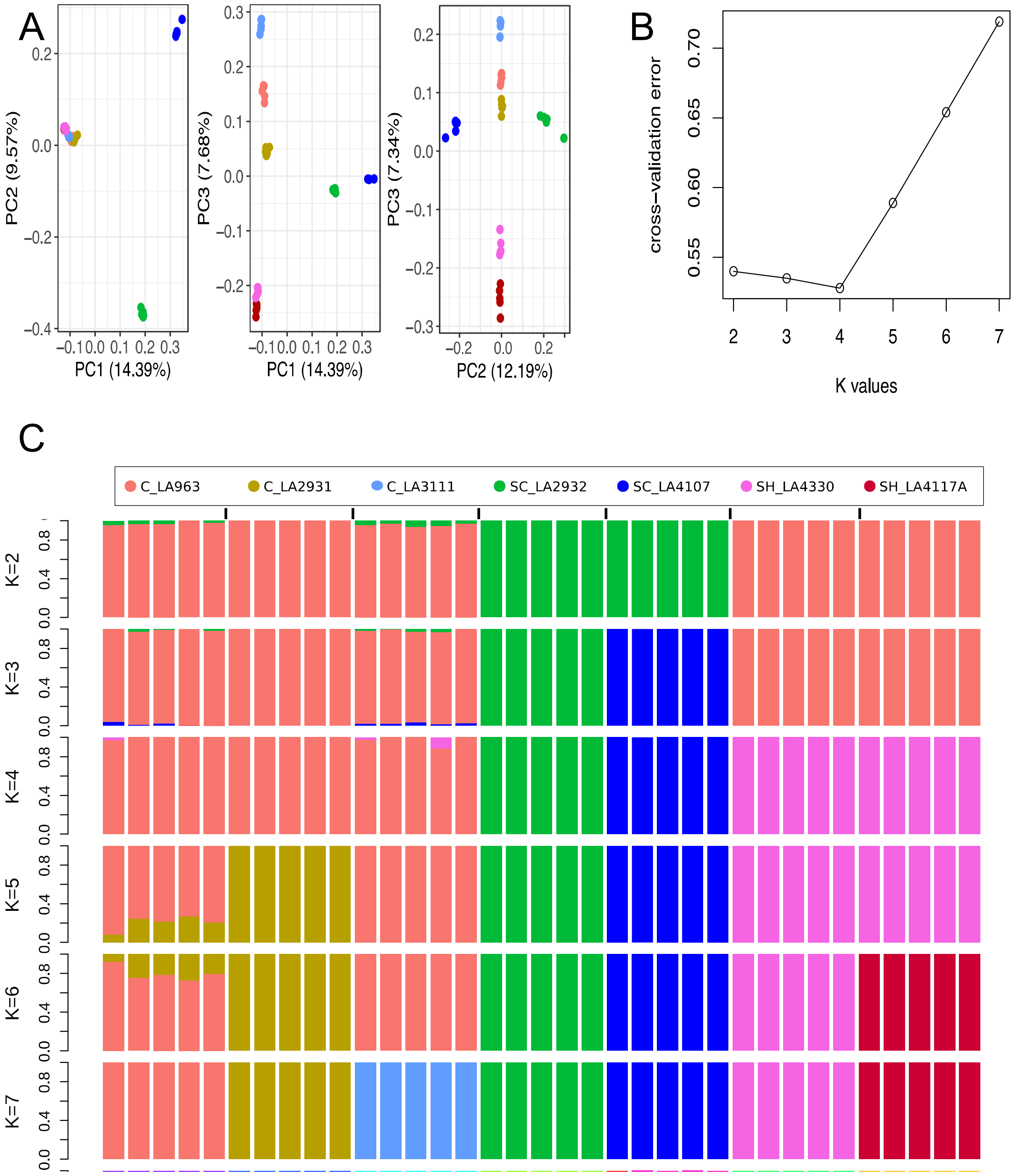


**Fig. S3.** The population structure analysis based on the whole-genome SNP data. (A) PCA based on the whole-genome SNP data. (B) the cross-validation error based on different K values. (C) The admixture based on the whole-genome SNP data.

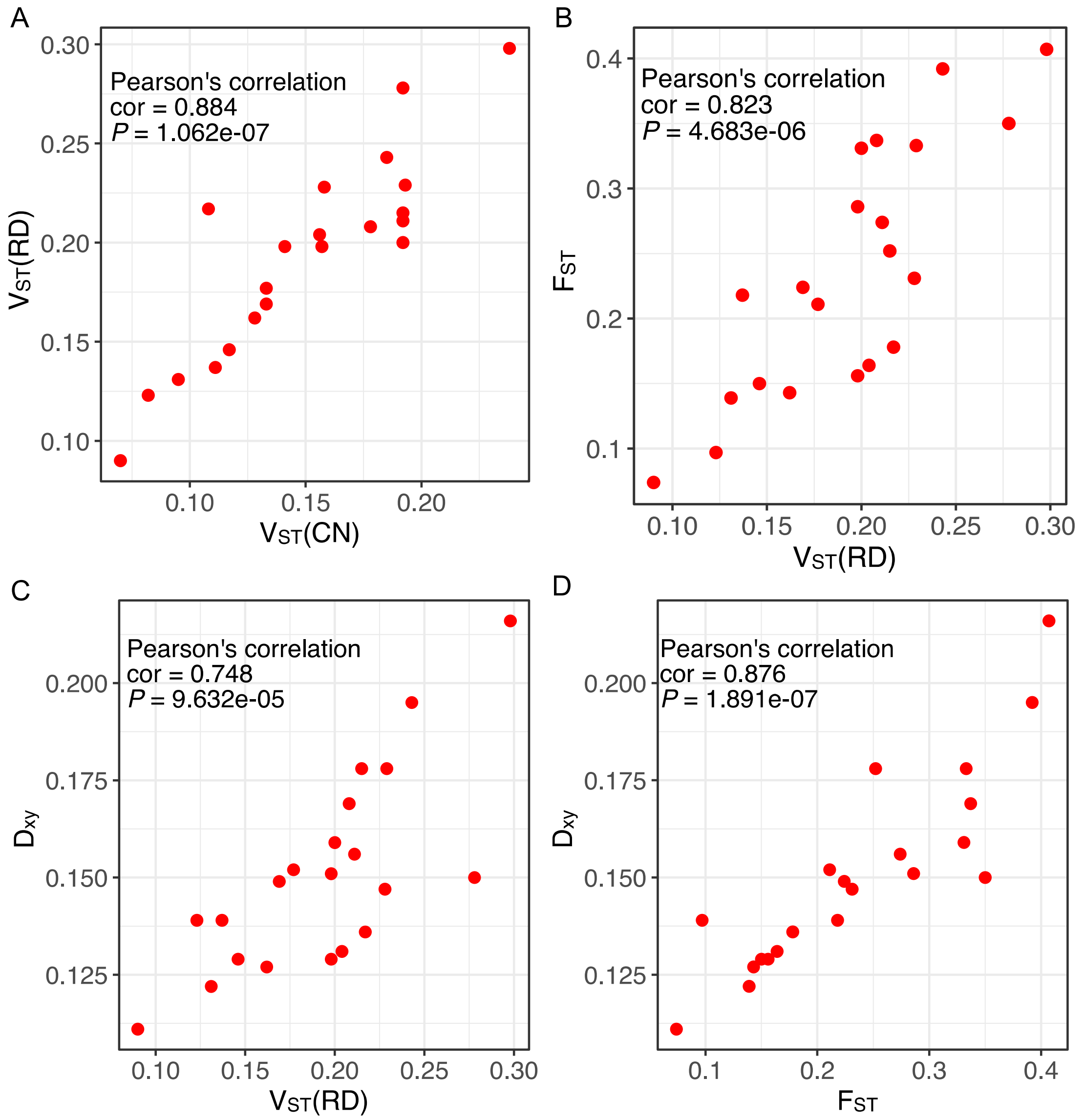


**Fig. S4.** The correlations between $V_{ST}$, $F_{ST}$ and $D_{xy}$ statistics. (A) The correlation between $V_{ST}$(CN) and $V_{ST}$(RD). (B) The correlation between $V_{ST}$(RD) and $F_{ST}$. (C) The correlation between $V_{ST}$(RD) and $D_{xy}$. (D) The correlation between $F_{ST}$ and $D_{xy}$.

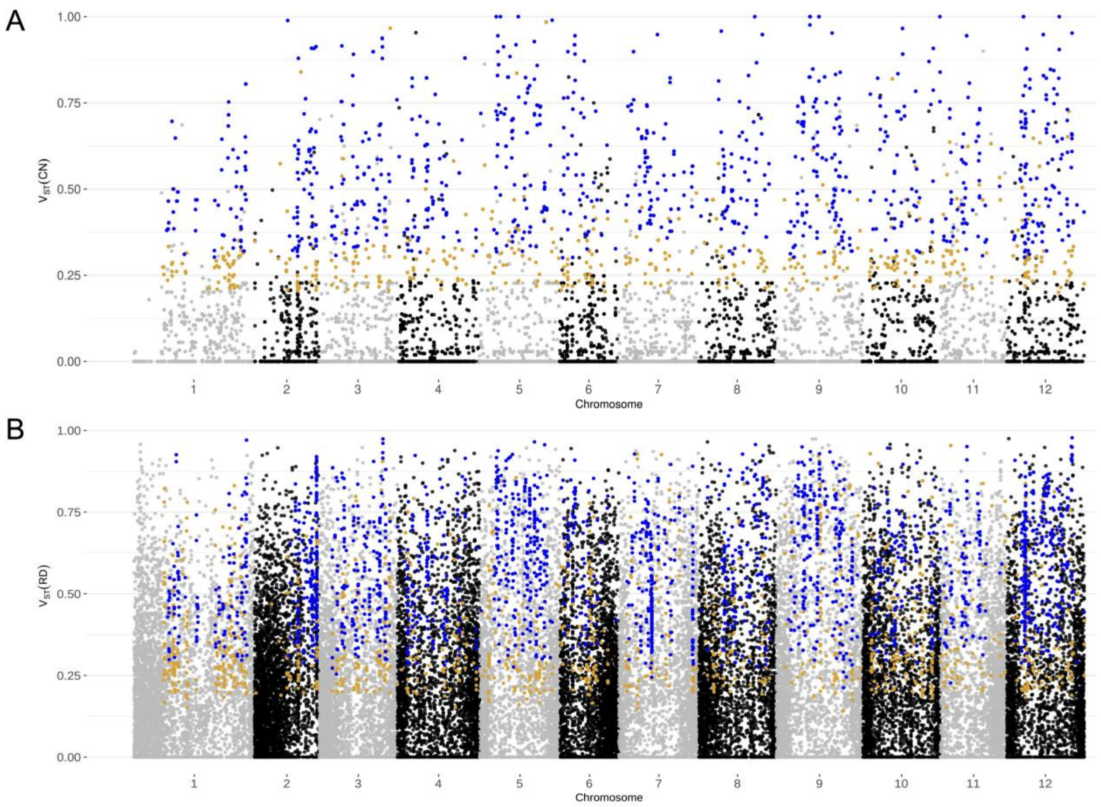


**Fig. S5.** The identification of the differentiated genes based on $V_{ST}(CN)$ (A) and $V_{ST}(RD)$ (B) using gene copy number (CN) quantified by Control-FREEC and Read Depth, respectively. The orange dots denote the CN-differentiated genes based on the 95$^{th}$ percentile from the distribution obtained with the permutation test (1,000 times), the blue dots denote the strongly CN-differentiated genes based on the 99$^{th}$ percentile from the distribution obtained with the permutation test (1,000 times). The dots represent genes.

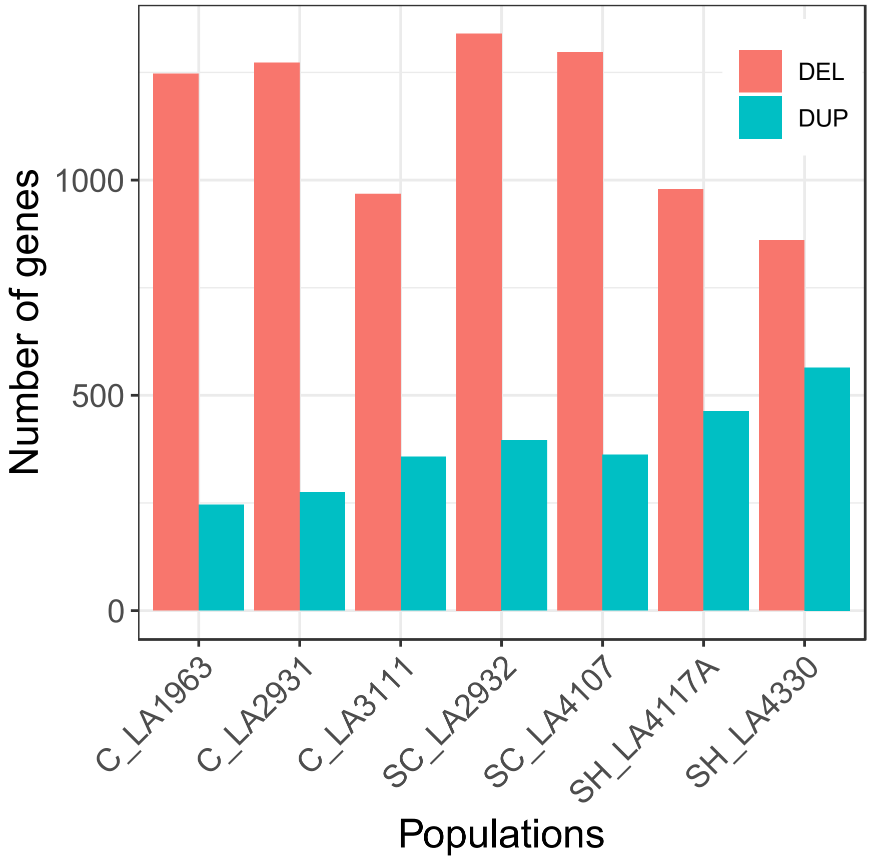


**Fig. S6.** The number of 3,539 CN-differentiated genes overlapping with deletion (DEL) and duplication (DUP) regions in seven populations.

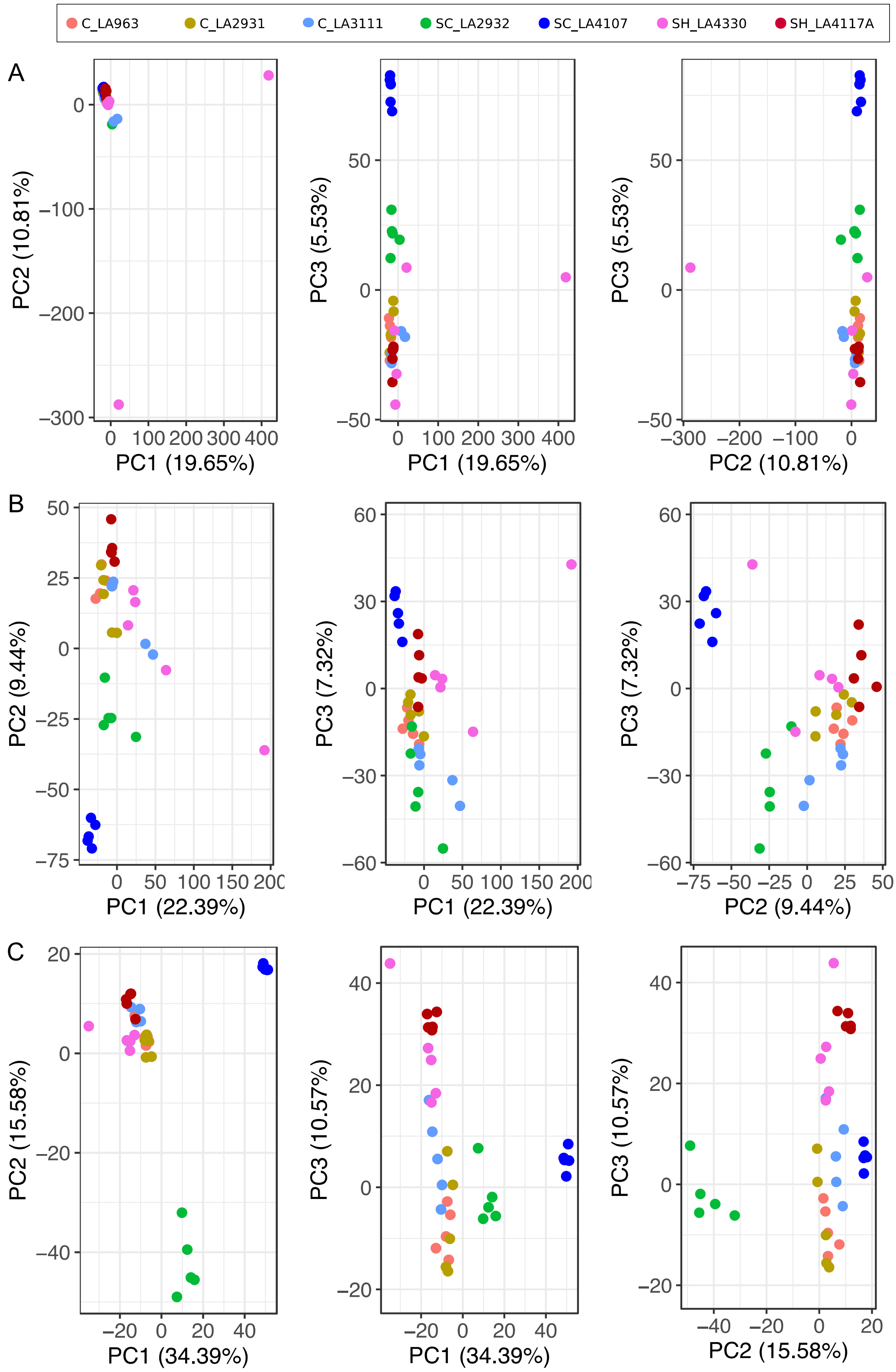


**Fig. S7.** PCA based on the gene copy number (CN) values of different gene sets in 35 individuals of seven populations. (A) PCA based on the CN values of 23,911 genes with mapped reads in 35 individuals. (B) PCA based on the CN values of 12,392 genes with $V_{ST}$(CN) > 0. (C) PCA based on the CN values of 2,192 strongly differentiated genes (see Table S5).

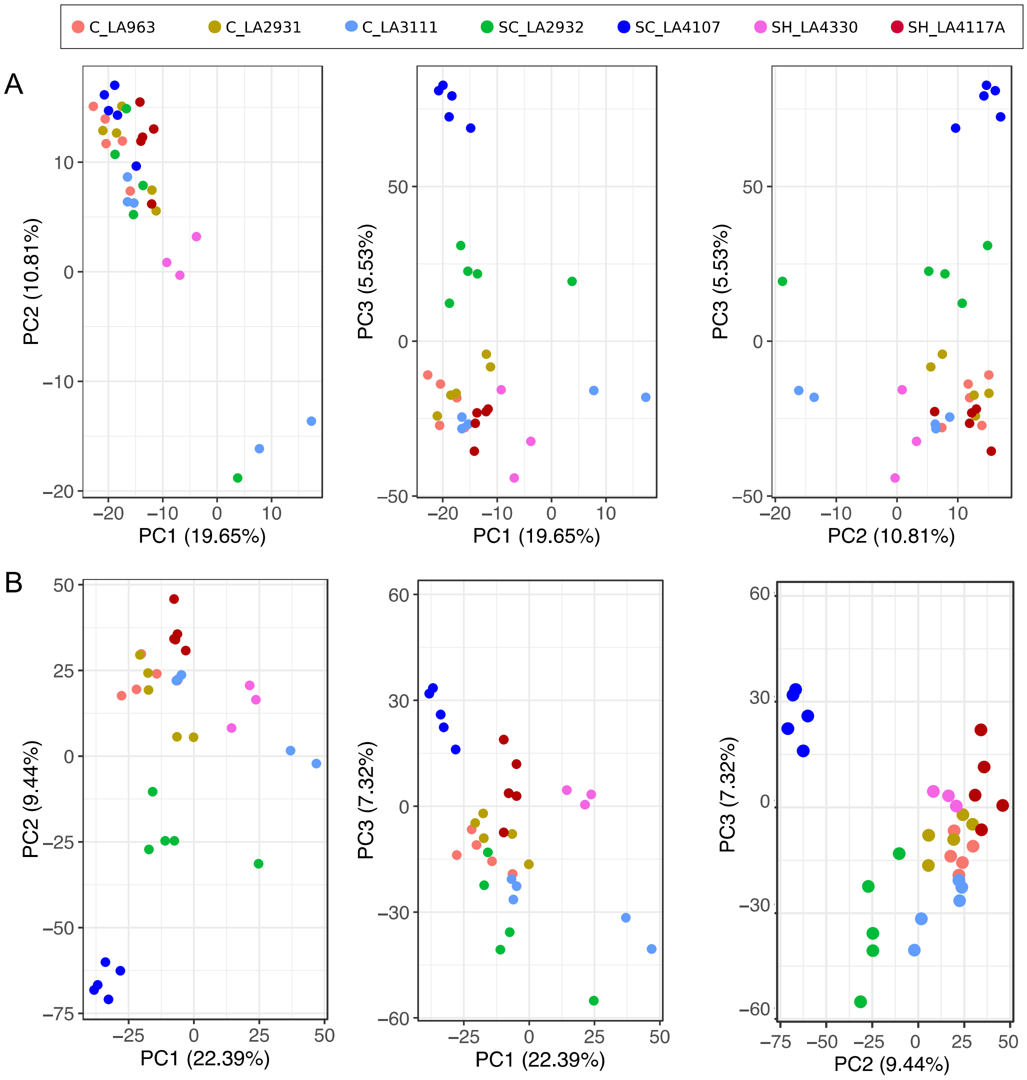


**Fig. S8.** Reconstructed PCA based on the gene copy number (CN) values of different gene sets in 35 individuals of seven populations removed the two outliers in Fig. S7A. The individuals in Fig. S7A and B were affected by two outliers causing most individuals to cluster together, so we redrew the PCA plots to remove the two outliers. (A) PCA based on the CN values of 23,911 genes with mapped reads in 35 individuals removed the two outliers in Fig. S7A. (B) PCA based on the CN values of 12,392 genes with $V_{ST}$(CN) > 0 removed the two outliers in Fig. S7A.

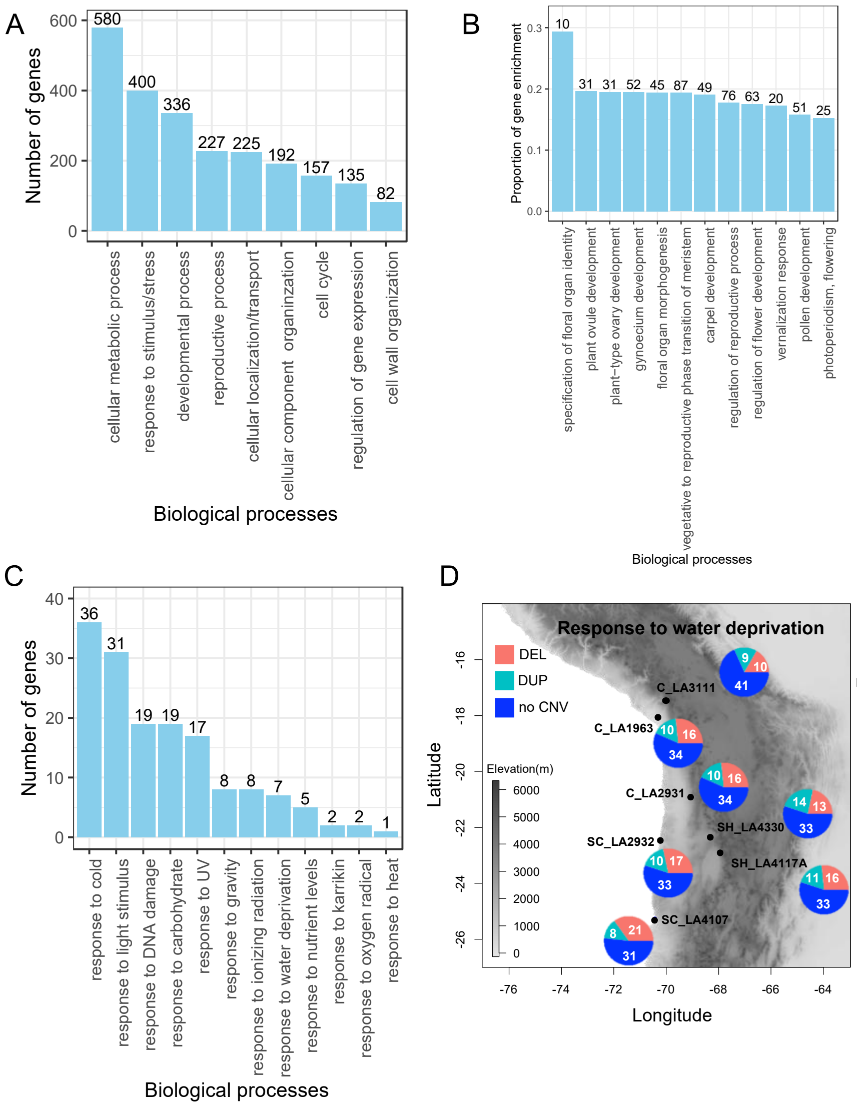


**Fig. S9.** GO enrichment analysis of 3,539 copy number (CN) differentiated genes. (A) The summary of GO enrichment. All significantly enriched GO terms ($P < 0.05$) were assigned into nine categories. (B) The proportions of differentiated genes enriched in different reproductive processes. The proportion of gene enrichment is equal to the number of genes enriched in one GO category divided by the number of background genes in this category. The number on top of each bar represents the number of genes enriched in that GO category. (C) The number of genes associated with the response to external stimulus/stresses overlapping with genes enriched in reproductive process. (D) the proportions of 60 CN-differentiated genes involved in response to water deprivation overlappping with deletion (DEL), duplication (DUP) or absence of CNV (no CNV) in the seven populations, respectively. The pie charts denote the proportions of CN-differentiated genes overlapped with DEL, DUP or absence of CNV (see also Table S6). The numbers in the pie chart indicate the number of genes overlapping with deletion, duplication or absence of CNV.

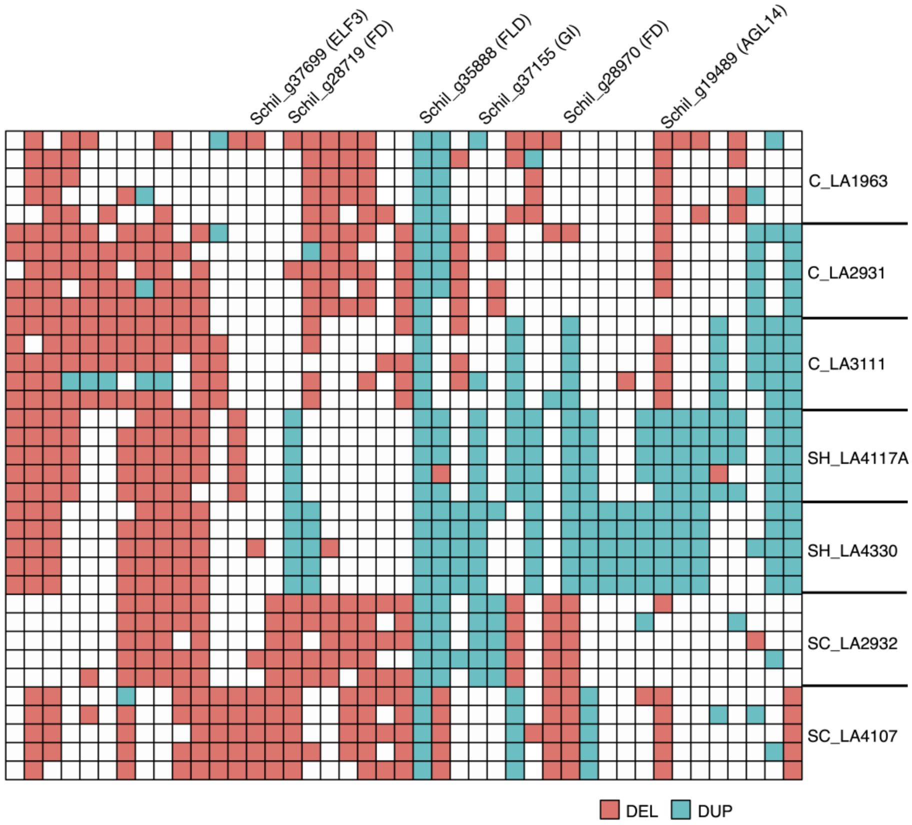


**Fig. S10.** CN-differentiated genes enriched in photoperiod and vernalization pathways show different CNV patterns across seven populations. Putative homologs (see also Dataset S5) are validated as flowering regulatory genes in other plant species named on the top of the matrix. White boxes indicate genes without CNV.

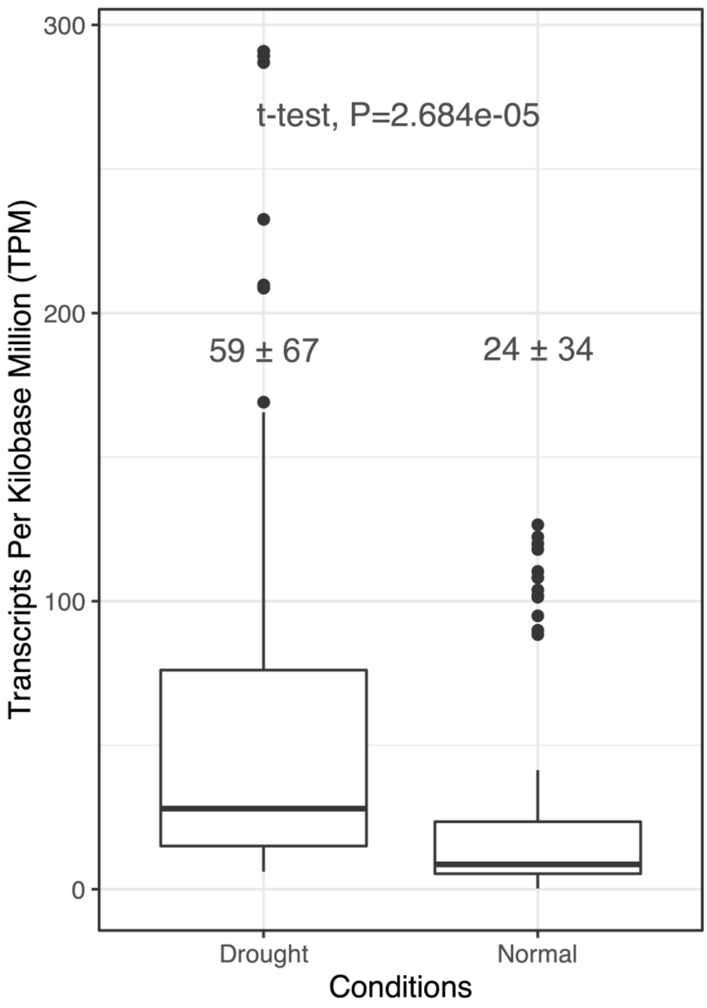


**Fig. S11.** Gene expression of 11 CN-differentiated genes associated with a response to water deprivation under drought and normal conditions. These 11 CN-differentiated genes in this study are overlapped with the drought-responsive genes identified from the previous comparative transcriptomic study (Wei et al. 2023a).

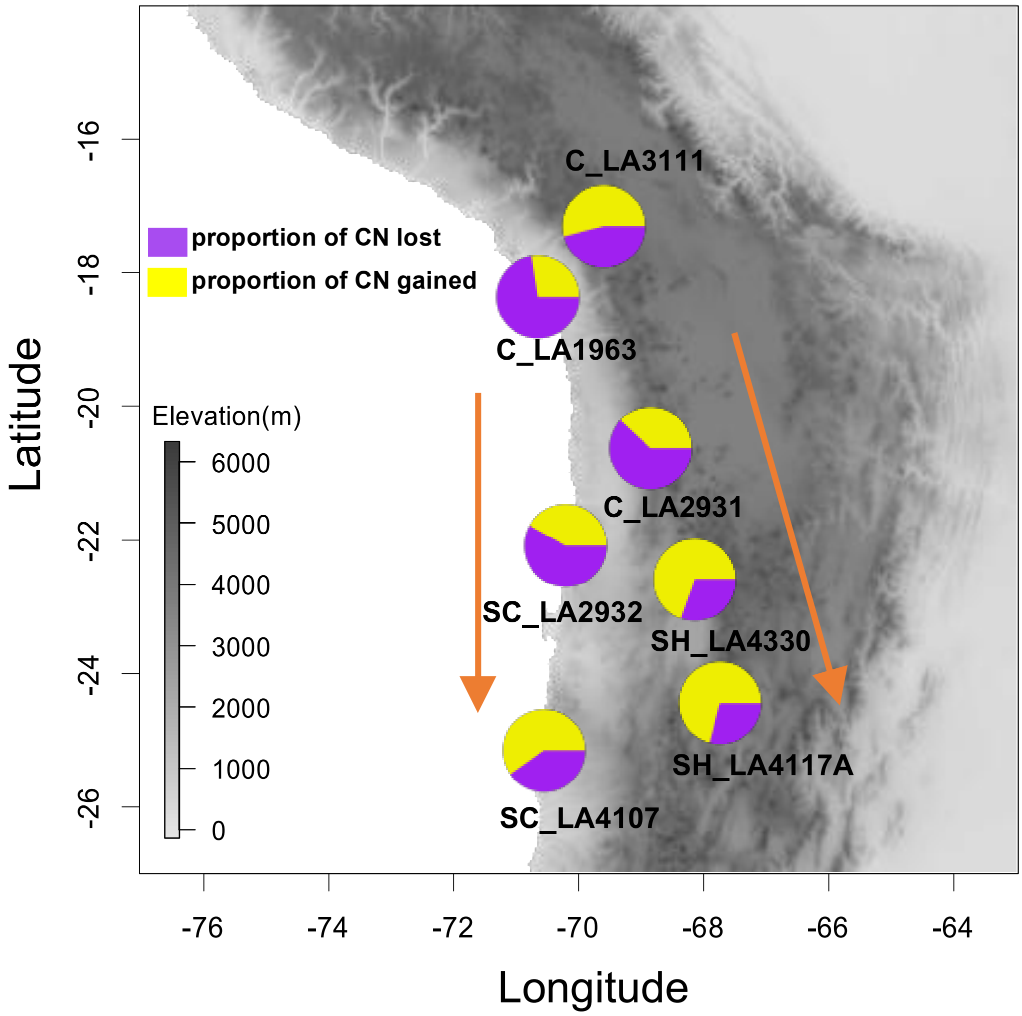


**Fig. S12.** The expansion and contraction of the copy number (CN) differentiated genes in different populations using the reference genome of *S. pennellii*. The map and pie charts show that the dynamics of copy number (CN) loss and gain in the processes of two southward colonization events, first to the southern coast (SC) and second to the southern highland (SH) (orange arrows) using the reference genome of *S. pennellii* (see also Table S9). The proportion of CN gains or losses for each population is defined as the number of CN gains or losses divided by the sum of the number of CN gains and losses.

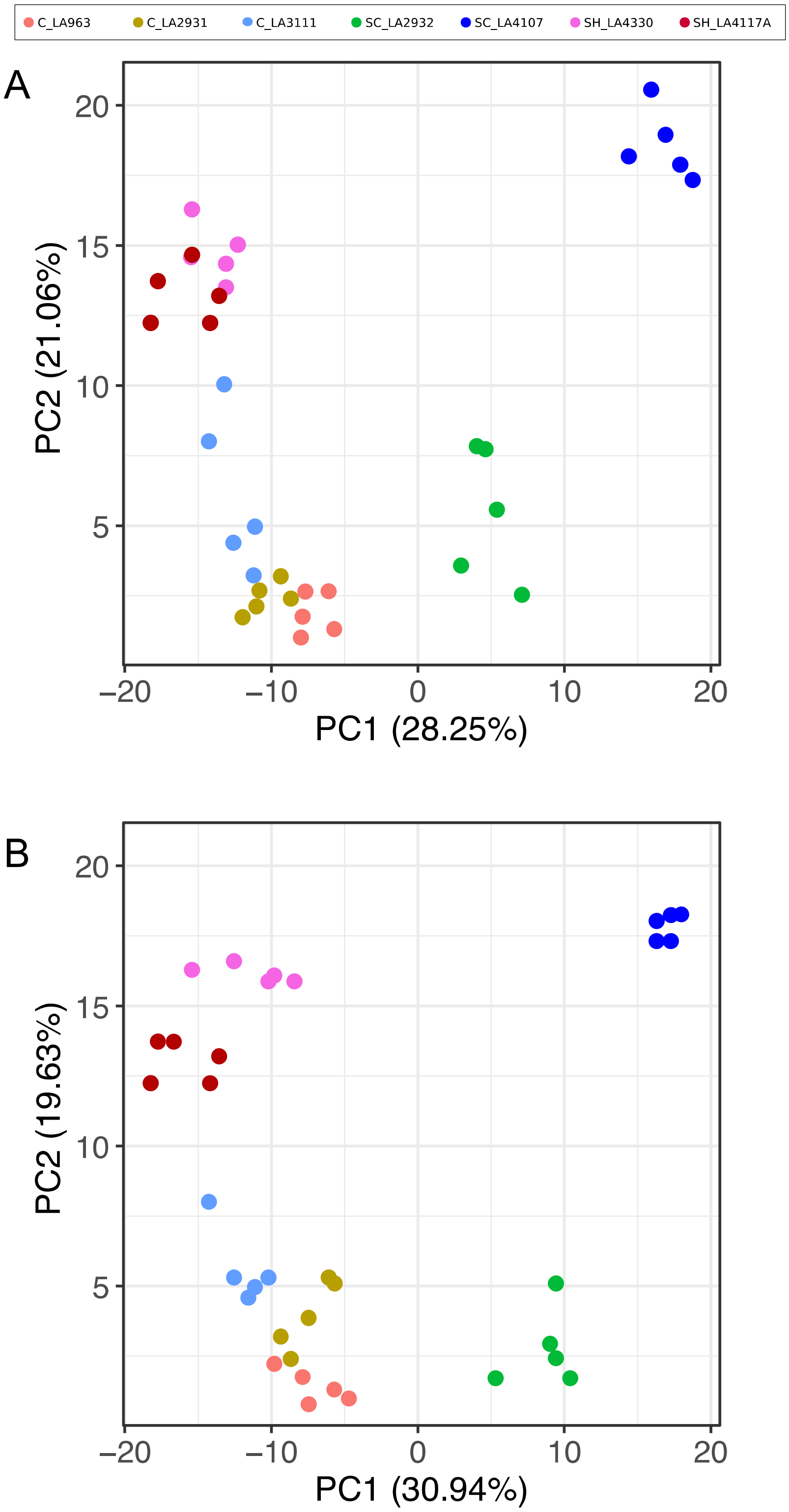


**Fig. S13.** PCA based on the copy number of rapidly evolving genes with significant CN expansion or contraction relative to the reference genome of *S. chilense* (A) and *S. pennellii* (B).

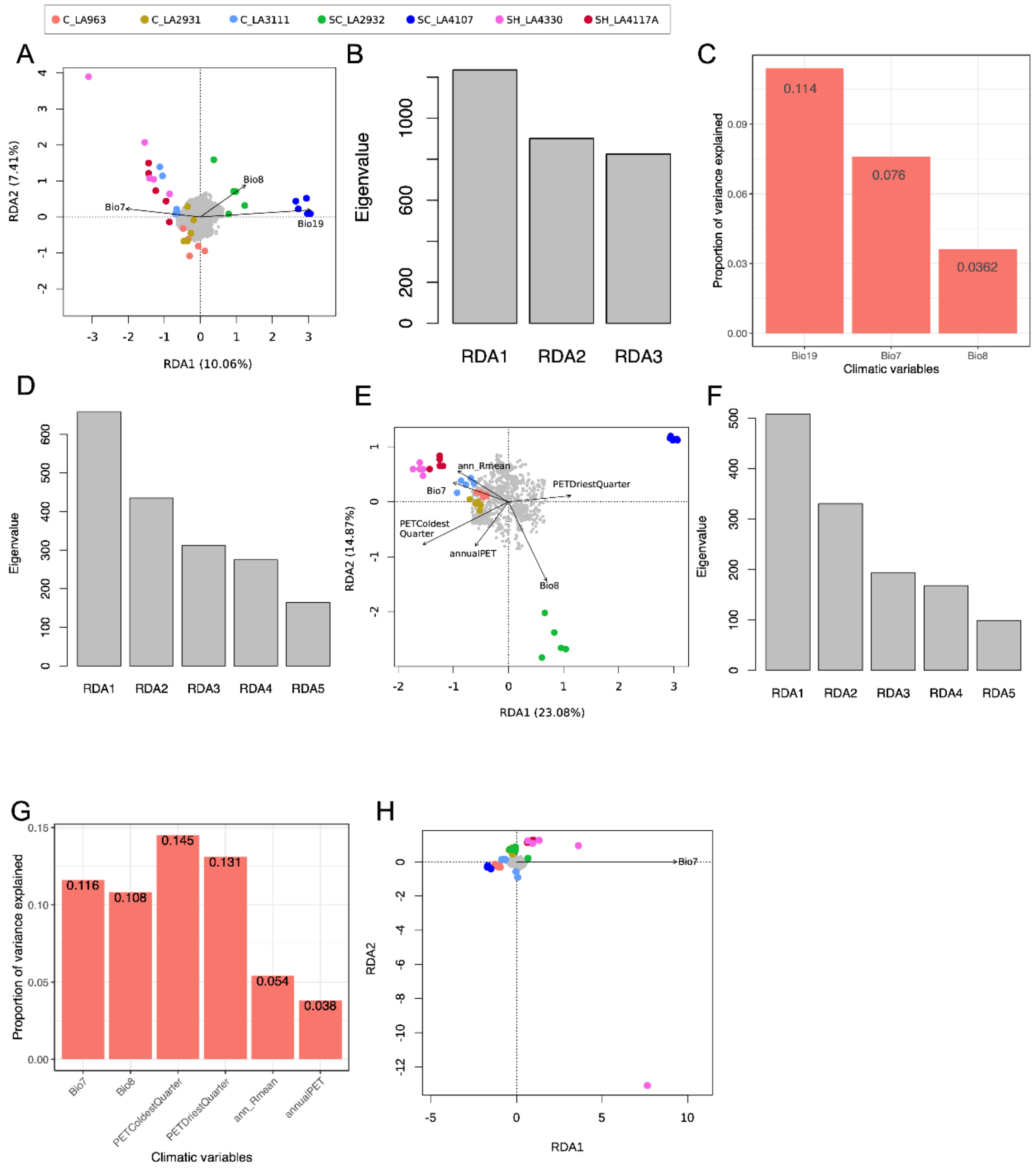


**Fig. S14.** Results of the redundancy analysis based on the climatic variables and gene copy number (CN) of the different gene sets. (A)The RDA model based on CN of 12,391 genes with $V_{ST} > 0$. (B) Eigenvalues of three significant ordination axes in an RDA model based on CN of 12,391 genes with $V_{ST} > 0$. (C) Proportion of variance explained by three overrepresented climate variables in an RDA model based on CN of 12,391 genes with $V_{ST} > 0$. (D) Eigenvalues of five significant ordination axes in an RDA model based on CN of 3,539 differentiated genes. (E) The RDA model based on CN of 2,192 strongly differentiated genes. (F) Eigenvalues of five significant ordination axes in an RDA model based on CN of 2,192 strongly differentiated genes. (G) Proportion of variance explained by six overrepresented climate variables in an RDA model based on CN of 2,192 strongly differentiated genes. (H) The RDA model shows no significant result based on CN of 20,372 non-CN differentiated genes. In the RDA models, arrows indicate the direction of the climatic variables associated with the different populations, and the projection of arrows onto an ordination axis show the correlation with that axis. The grey points denote the CN-differentiated genes. The dots represent individual plants and are colored by population. C: central; SH: southern highland; SC: southern coast.

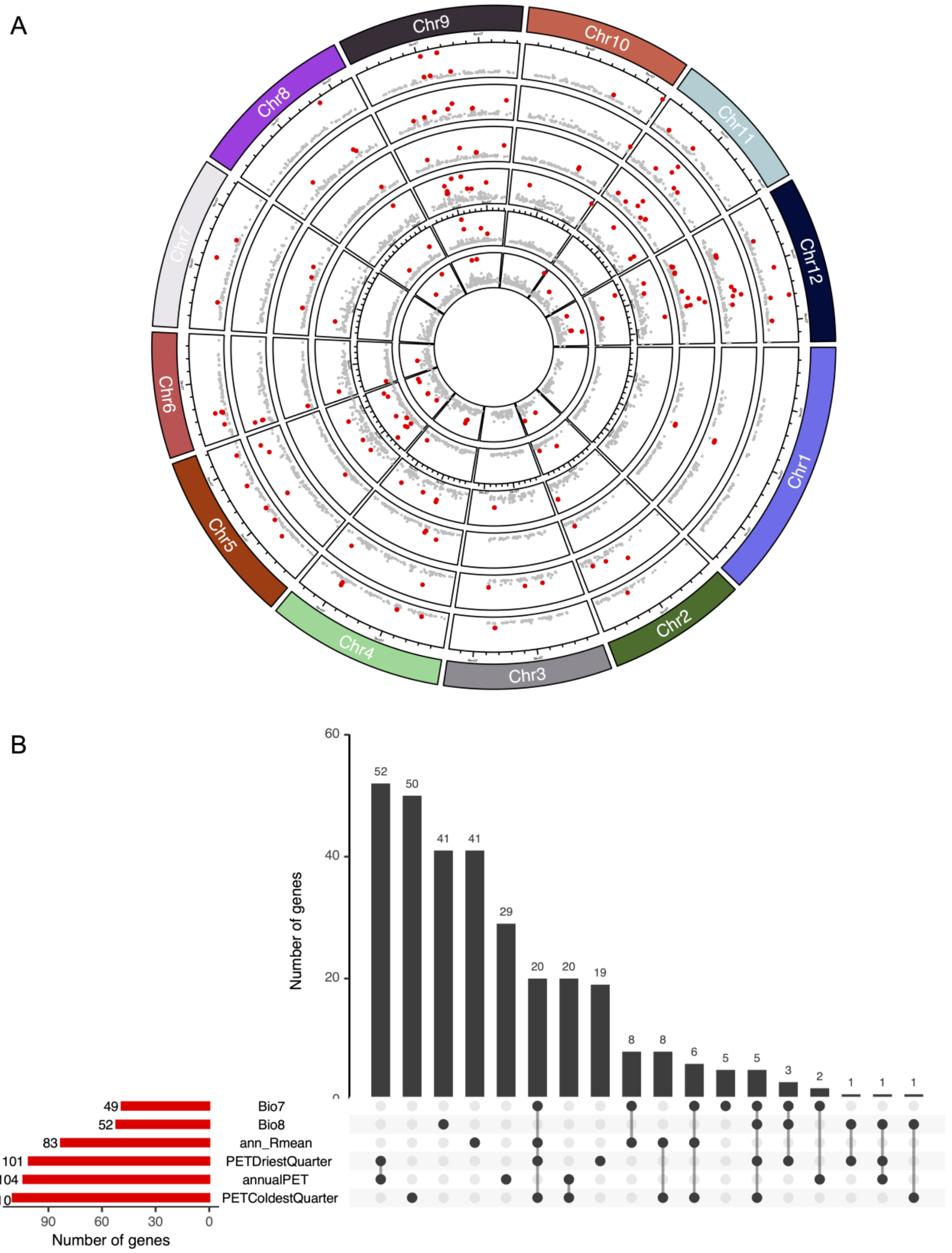


**Fig. S15.** The genome-environment association (GEA) analysis between gene copy number (CN) and the climatic variables using LFMM2. (A) Strength of evidence for an association between gene CN and six climatic variables based on the RDA. From the center to the margin, the results are shown for Bio7, Bio8, ann_Rmean, PETDriestQuarter, annualPET, and PETColdestQuarter. The horizontal axis denotes the position of the gene on the chromosome and the vertical axis denotes the values of $-\log_{10}(P)$. Red dots represent candidate genes significantly associated with specific climate variables. (B) The number of candidate genes associated with six climatic variables (red bar), respectively, and shared candidate genes across climatic variables (black bar).

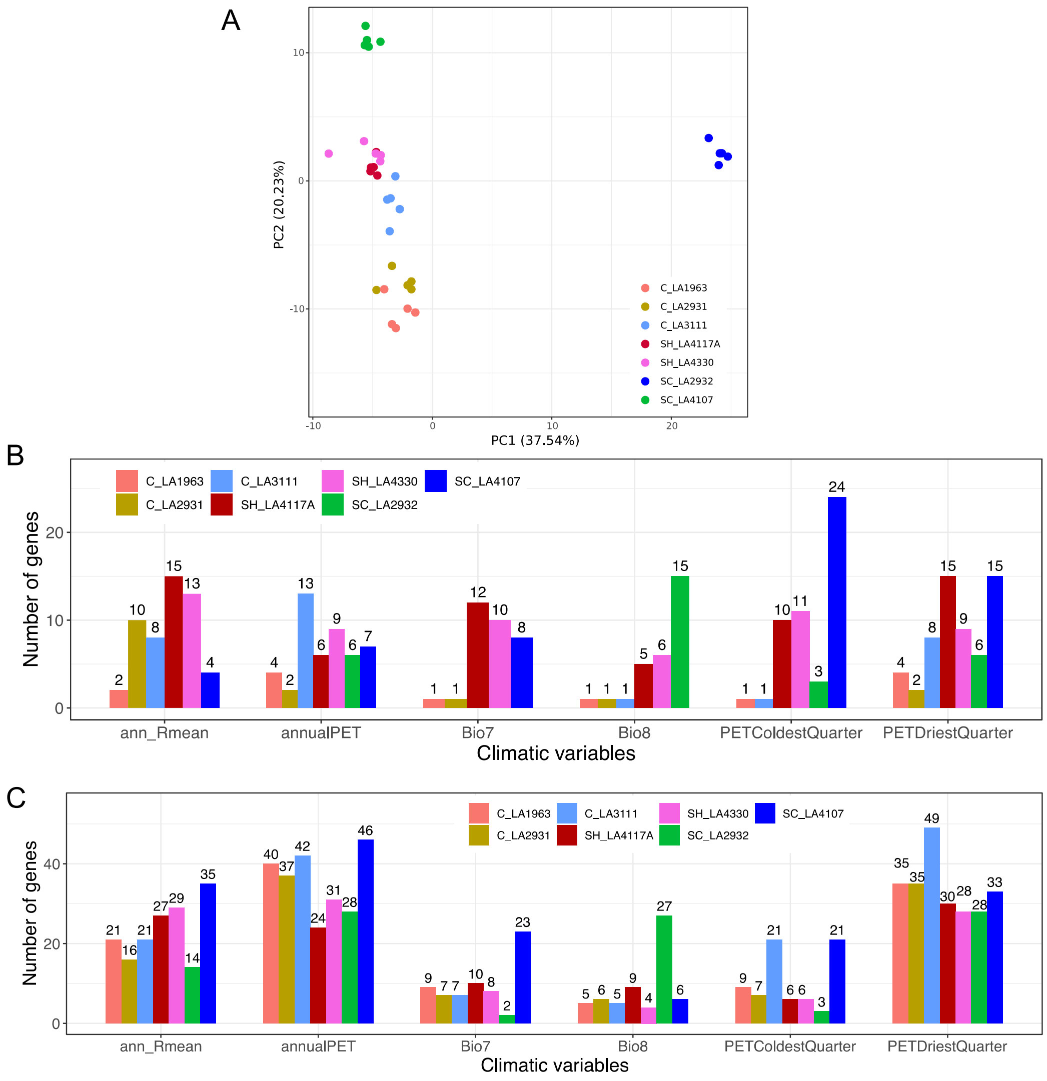


**Fig. S16.** Summary of candidate genes associated with the climatic variables. (A) PCA based on the copy number (CN) of 312 candidate genes identified by LFMM2 as being associated with six climatic variables. (B) The number of candidate genes identified by LFMM2 located at duplication (DUP) regions in seven populations. (C) The number of candidate genes identified by LFMM2 located at deletion (DEL) regions in seven populations.

**Table S1.** The number of deletion (DEL) and duplication (DUP) in each population

| Populations | DEL | DUP |
|---|---|---|
| C_LA1963 | 68393 | 26228 |
| C_LA2931 | 66859 | 25970 |
| C_LA3111 | 58859 | 25133 |
| SC_LA2932 | 56914 | 22576 |
| SC_LA4107 | 51849 | 21165 |
| SH_LA4117A | 54314 | 22504 |
| SH_LA4330 | 55661 | 23117 |

**Table S2**. The number of deletion (DEL) and duplication (DUP) in each individual.

| Groups | Populations | Individuals | DEL | DUP | Total |
|---|---|---|---|---|---|
| central | C_LA1963 | C_LA1963_t1 | 31891 | 11736 | 43627 |
| central | C_LA1963 | C_LA1963_t2 | 28596 | 10605 | 39201 |
| central | C_LA1963 | C_LA1963_t5 | 31627 | 11908 | 43535 |
| central | C_LA1963 | C_LA1963_t7 | 29069 | 10913 | 39982 |
| central | C_LA1963 | C_LA1963_t9 | 31312 | 11679 | 42991 |
| central | C_LA2931 | C_LA2931_t2 | 30586 | 11342 | 41928 |
| central | C_LA2931 | C_LA2931_t3 | 30871 | 11334 | 42205 |
| central | C_LA2931 | C_LA2931_t4 | 31676 | 11809 | 43485 |
| central | C_LA2931 | C_LA2931_t5 | 32074 | 11730 | 43804 |
| central | C_LA2931 | C_LA2931_t6 | 31133 | 11556 | 42689 |
| central | C_LA3111 | C_LA3111_t3 | 28357 | 11789 | 40146 |
| central | C_LA3111 | C_LA3111_t5 | 30898 | 12507 | 43405 |
| central | C_LA3111 | C_LA3111_t9 | 25406 | 10597 | 36003 |
| central | C_LA3111 | C_LA3111_t10 | 27963 | 11594 | 39557 |
| central | C_LA3111 | C_LA3111_t15 | 27623 | 11765 | 39388 |
| southern coast | SC_LA2932 | SC_LA2932_1 | 34249 | 12330 | 46579 |
| southern coast | SC_LA2932 | SC_LA2932_8 | 31684 | 11379 | 43063 |
| southern coast | SC_LA2932 | SC_LA2932_t2 | 32097 | 11514 | 43611 |
| southern coast | SC_LA2932 | SC_LA2932_20 | 31230 | 11074 | 42304 |
| southern coast | SC_LA2932 | SC_LA2932_22 | 30155 | 10765 | 40920 |
| southern coast | SC_LA4107 | SC_LA4107_3 | 30991 | 11103 | 42094 |
| southern coast | SC_LA4107 | SC_LA4107_6 | 33141 | 12096 | 45237 |
| southern coast | SC_LA4107 | SC_LA4107_9 | 32849 | 11761 | 44610 |
| southern coast | SC_LA4107 | SC_LA4107_t5 | 31243 | 11456 | 42699 |
| southern coast | SC_LA4107 | SC_LA4107_t11 | 30818 | 11006 | 41824 |
| southern highland | SH_LA4117A | SH_LA4117A_1 | 27913 | 9391 | 37304 |
| southern highland | SH_LA4117A | SH_LA4117A_4 | 28128 | 10452 | 38580 |
| southern highland | SH_LA4117A | SH_LA4117A_5 | 28858 | 9865 | 38723 |
| southern highland | SH_LA4117A | SH_LA4117A_10 | 27772 | 10131 | 37903 |
| southern highland | SH_LA4117A | SH_LA4117A_15 | 24905 | 7018 | 31923 |
| southern highland | SH_LA4330 | SH_LA4330_t1 | 29186 | 11019 | 40205 |
| southern highland | SH_LA4330 | SH_LA4330_t4 | 32217 | 12281 | 44498 |
| southern highland | SH_LA4330 | SH_LA4330_t6 | 26220 | 9823 | 36043 |
| southern highland | SH_LA4330 | SH_LA4330_t9 | 31562 | 11964 | 43526 |
| southern highland | SH_LA4330 | SH_LA4330_t12 | 30934 | 11610 | 42544 |

**Table S3.** The validation of pipeline of CNV calling using 1,000 simulated deletions (DELs) and duplications (DUPs), respectively.

| CNV caller | Number of DEL | Number of DUP |
|---|---|---|
| Lumpy | 878 (21) | 842(13) |
| Manta | 795 (29) | 774(36) |
| Wham | 767 (33) | 698 (19) |
| Delly | 849 (40) | 861 (22) |
| SURVIVOR merged | 918(12) | 879 (4) |

Note: The numbers given outside and inside of the parentheses represent the CNV number of true and false positives called using the pipeline in this study based on simulated data.

**Table S4.** Measures of population differentiation based on copy number ($V_{ST}$(RD) and $V_{ST}$(CN)) and SNPs ($F_{ST}$).

| Pairwise populations | $V_{ST}$(RD) | $V_{ST}$(CN) | $F_{ST}$ | *D*xy |
|---|---|---|---|---|
| C_LA1963 vs C_LA2931 | 0.090 ±0.153 | 0.070±0.125 | 0.074±0.131 | 0.111±0.050 |
| C_LA1963 vs C_LA3111 | 0.123±0.177 | 0.082±0.138 | 0.097±0.146 | 0.139±0.051 |
| C_LA2931 vs C_LA3111 | 0.131±0.194 | 0.095±0.162 | 0.139±0.155 | 0.122±0.050 |
| SH_LA4117A vs SH_LA4330 | 0.217±0.239 | 0.108±0.188 | 0.178±0.247 | 0.136±0.054 |
| C_LA3111 vs SH_LA4330 | 0.137±0.212 | 0.111±0.177 | 0.218±0.227 | 0.139±0.056 |
| C_LA2931 vs SH_LA4330 | 0.146±0.198 | 0.117±0.165 | 0.150±0.163 | 0.129±0.055 |
| C_LA1963 vs SH_LA4330 | 0.162±0.203 | 0.128±0.190 | 0.143±0.147 | 0.127±0.055 |
| C_LA2931 vs SC_LA2932 | 0.169±0.245 | 0.133±0.134 | 0.224±0.270 | 0.149±0.062 |
| C_LA1963 vs SC_LA2932 | 0.177±0.248 | 0.133±0.174 | 0.211±0.196 | 0.152±0.064 |
| C_LA1963 vs SH_LA4117A | 0.198±0.244 | 0.141±0.189 | 0.156±0.149 | 0.129±0.055 |
| C_LA2931 vs SH_LA4117A | 0.204±0.248 | 0.156±0.206 | 0.164±0.172 | 0.131±0.055 |
| SC_LA2932 vs C_LA3111 | 0.198±0.276 | 0.157±0.196 | 0.286±0.311 | 0.151±0.067 |
| C_LA3111 vs SH_LA4117A | 0.228±0.257 | 0.158±0.218 | 0.231±0.255 | 0.147±0.057 |
| SC_LA2932 vs SH_LA4330 | 0.208±0.267 | 0.178±0.230 | 0.337±0.382 | 0.169±0.065 |
| SC_LA4107 vs SH_LA4330 | 0.243±0.291 | 0.185±0.259 | 0.392±0.431 | 0.195±0.067 |
| C_LA1963 vs SC_LA4107 | 0.215±0.272 | 0.192±0.236 | 0.252±0.277 | 0.178±0.067 |
| C_LA2931 vs SC_LA4107 | 0.211±0.275 | 0.192±0.244 | 0.274±0.278 | 0.156±0.064 |
| SC_LA2932 vs SC_LA4107 | 0.200±0.281 | 0.192±0.254 | 0.331±0.337 | 0.159±0.070 |
| SC_LA2932 vs SH_LA4117A | 0.278±0.300 | 0.192±0.259 | 0.350±0.414 | 0.150±0.064 |
| C_LA3111 vs SC_LA4107 | 0.229±0.284 | 0.193±0.262 | 0.333±0.401 | 0.178±0.070 |
| SC_LA4107 vs SH_LA4117A | 0.298±0.315 | 0.238±0.287 | 0.407±0.448 | 0.216±0.067 |

**Table S5.** Number of candidate genes with differentiated gene CN across seven populations

| | Differentiated (95$^{th}$ percentile) | | Strongly differentiated (99$^{th}$ percentile) | |
|---|---|---|---|---|
| $V_{ST}$ | Threshold | Number of genes | Threshold | Number of genes |
| $V_{ST}$(CN) | 0.194 | 4,843 | 0.305 | 3,219 |
| $V_{ST}$(RD) | 0.157 | 16,655 | 0.244 | 12,228 |
| Overlaps | | 3,539 | | 2,192 |

**Table S6.** Number of CN differentiated genes involved in four GO terms in seven populations located in deletion (DEL) and duplication (DUP) regions.

| Populations | photoperiod | | | vernalization | | | response to water deprivation | | | root development | | |
|---|---|---|---|---|---|---|---|---|---|---|---|---|
| | DEL | DUP | no CNV | DEL | DUP | no CNV | DEL | DUP | no CNV | DEL | DUP | no CNV |
| C_LA1963 | 6 | 1 | 18 | 8 | 1 | 11 | 16 | 10 | 34 | 26 | 17 | 30 |
| C_LA2931 | 10 | 1 | 14 | 9 | 3 | 8 | 16 | 10 | 34 | 12 | 4 | 57 |
| C_LA3111 | 8 | 2 | 15 | 7 | 5 | 8 | 10 | 9 | 41 | 11 | 8 | 54 |
| SC_LA2932 | 10 | 2 | 13 | 7 | 2 | 11 | 17 | 10 | 33 | 27 | 18 | 28 |
| SC_LA4107 | 9 | 2 | 14 | 10 | 1 | 9 | 21 | 8 | 31 | 25 | 18 | 30 |
| SH_LA4117A | 5 | 8 | 12 | 5 | 8 | 7 | 16 | 11 | 33 | 14 | 7 | 52 |
| SH_LA4330 | 4 | 10 | 11 | 4 | 7 | 9 | 13 | 14 | 33 | 12 | 6 | 55 |

**Table S7.** Summary of CN expansion and contraction in different branches or populations using the reference of *S. pennellii*.

| Groups or Populations | Number of CN expanded genes | Number of CN contracted genes | Number of CN gained | number of CN contracted | [a]Rate of average expansion or contraction | [b]Number of rapidly evolving genes |
|---|---|---|---|---|---|---|
| inland | 63 | 48 | 344 | 86 | 2.324 | 28(+21/-7) |
| Central (C) | 186 | 568 | 427 | 937 | -0.676 | 35(+15/-20) |
| southern highland (SH) | 522 | 384 | 1506 | 648 | 0.947 | 51(+38/-13) |
| southern coast (SC) | 67 | 115 | 184 | 211 | -0.1483 | 11(+4/-7) |
| C_LA1963 | 149 | 456 | 322 | 851 | -0.874 | 27(+9/-18) |
| C_LA2931 | 116 | 292 | 360 | 587 | -0.556 | 14(+3/-11) |
| C_LA3111 | 215 | 374 | 762 | 646 | 0.197 | 19(+11/-8) |
| SH_LA4117A | 627 | 484 | 2007 | 824 | 1.065 | 54(+39/-15) |
| SH_LA4330 | 471 | 304 | 1340 | 588 | 0.970 | 43(+43/-0) |
| SC_LA2932 | 215 | 559 | 747 | 1035 | -0.372 | 17(+5/-12) |
| SC_LA4107 | 384 | 306 | 1132 | 692 | 0.638 | 38(+26/-12) |

[a]Rate of average expansion or contraction = (Number of CN gains - Number of CN losses) / (Number of CN expanded genes + Number of CN contracted genes). Positive values indicate CN expansion and negative values indicate CN contraction.

[b]The rapidly evolving genes indicate significant higher CN expansion or contraction ($P$ values were obtained by the Viterbi method with the randomly generated likelihood distribution; $P < 0.05$) across the different groups or populations. Values outside parentheses represent the total number of the rapidly evolving genes. Positive values in parentheses denote the number of significantly expanded genes and negative values denote the number of significantly contracted genes.

**Table S8.** Number of CN gains (positive values) and CN losses (negative values) for rapidly evolving genes related to photosynthesis in different subgroups representing four different habitats.

| Genes | Central | Southern highland | SC_LA2932 | SC_LA4107 |
|---|---|---|---|---|
| Schil_g10783 | - | 19 | - | - |
| Schil_g12727 | - | - | - | 15 |
| Schil_g13768 | - | - | -6 | - |
| Schil_g16334 | -20 | 25 | -21 | 21 |
| Schil_g16755 | - | 9 | - | - |
| Schil_g17486 | -5 | - | - | - |
| Schil_g17837 | -18 | 19 | -17 | 21 |
| Schil_g17839 | -18 | 19 | -17 | 21 |
| Schil_g17842 | -18 | 19 | -17 | 21 |
| Schil_g18042 | - | - | - | 16 |
| Schil_g19313 | -5 | 16 | -7 | - |
| Schil_g28833 | - | 9 | - | - |
| Schil_g29254 | -10 | 16 | -11 | 10 |
| Schil_g34328 | -14 | 14 | -17 | 20 |
| Schil_g34329 | -14 | 14 | -17 | 20 |
| Schil_g38065 | - | 7 | - | - |

**Table S9.** Number of CN-differentiated genes associated with temperature annual range (Bio7) and solar radiation (ann_Rmean) in seven populations located in deletion (DEL) and duplication (DUP) regions.

| Populations | DEL | DUP | no CNV |
|---|---|---|---|
| C_LA1963 | 7 | 1 | 26 |
| C_LA2931 | 6 | 3 | 25 |
| C_LA3111 | 2 | 1 | 31 |
| SC_LA2932 | 7 | 2 | 25 |
| SC_LA4107 | 10 | 1 | 23 |
| SH_LA4117A | 2 | 5 | 27 |
| SH_LA4330 | 6 | 6 | 22 |
